# Recovering directional brain networks under temporal undersampling: an application to schizophrenia

**DOI:** 10.64898/2026.08.09.743708

**Authors:** Mohammadsajad Abavisani, Kseniya Solovyeva, David Danks, Godfrey Pearlson, Vince Calhoun, Sergey Plis

## Abstract

Two decades of functional connectivity research have established schizophrenia as a disorder of distributed dysconnectivity, with a robust thalamocortical signature: reduced prefrontal and increased sensory coupling. A fundamental issue is that functional connectivity is undirected, operates at a single slow timescale, and cannot reveal causal direction. Moreover, the mismatch between BOLD sampling speed and neural dynamics can hide edges, fabricate spurious ones, and reverse the apparent orientation of causal relationships. To overcome these limitations, we introduce a general framework for estimating directed causal graphs from fMRI that explicitly accounts for temporal undersampling. We apply RnR, a causal discovery method built on the rate-agnostic RASL framework, to resting-state fMRI from the multi-site FBIRN cohort. Rather than returning a single directed graph, RnR recovers an equivalence class of directed graphs consistent with the observed data, each annotated with the sampling rate that would produce it and classifies each estimated orientation by its stability across inferred rates. This provides a principled basis for distinguishing directed interpretations that are safe to trust from those that are timescale-contingent. Benchmarking against five single-timescale estimators on schizophrenia data, RnR recovered substantially more group-differentiating directed edges, reproducing the field’s most replicated finding in directed form: a sensory-to-visual hyperconnectivity hub oriented from the post-central gyrus component to primary visual cortex. Through simulation, we show that coarse spatial resolution has shielded single-timescale methods from the full effects of undersampling, whereas finer parcellations will require explicit undersampling modeling. This work reframes fMRI effective connectivity estimation by treating undersampling as a fundamental property of the measurement, enabling directed interpretations that are grounded in the data-generating process.

## 1 Introduction

The idea that schizophrenia is a disorder of disrupted communication between brain regions is one of the most enduring in the field, and functional MRI has repeatedly localised this disruption to circuits linking the thalamus, sensory cortex, and prefrontal cortex. However, the correlation-based measures underpinning these findings share two fundamental limitations: they do not indicate the direction of influence between regions, and they are derived from a signal sampled considerably more slowly than the neural activity it reflects, a mismatch that can distort or even reverse apparent connectivity. Here we reanalyze a multi-site schizophrenia dataset using a causal-discovery method that directly models this slow-sampling problem and returns directed connections. The method reproduces the established connectivity differences between patients and controls, now with directional information attached, and uniquely identifies which of those directed edges are stable across sampling rates and which are not, reporting only the stable subset. We further demonstrate why measuring the brain at the coarse scale of regions, rather than neurons, has permitted slower, direction-blind methods to succeed thus far, and why moving to finer spatial resolution will render explicit modelling of the sampling rate essential.

Schizophrenia has long been viewed not as a disease of any single brain region but as a disorder of connections, and more specifically, a failure of functional integration among distributed neural systems. This “dysconnection” framing, given its modern form by **(author?)** [18], holds that the core symptoms arise from aberrant coupling between brain regions rather than from focal lesions [34]. Two decades of functional neuroimaging have consistently confirmed this view: the most reproducible findings in schizophrenia (across seed-based, independent-component, dynamic, and graph-theoretic analyses) are distributed, network-level, and centered on cortico-subcortical hubs.

Among the most replicated findings is the thalamocortical signature: reduced coupling between the thalamus and prefrontal cortex alongside increased coupling between the thalamus and sensorimotor, auditory and visual cortices [45, 13, 15, 5]. At the level of canonical cortical networks the recurring picture is within-network hypoconnectivity in default-mode and self-referential systems, disturbed coupling of the salience network with default-mode and executive systems, and a broad reduction in segregation between task-positive and task-negative systems. Both patterns are read through one substantive hypothesis: a thalamus that transmits insufficiently filtered sensory input to cortex under weakened prefrontal regulation [4]. The next section (Related Works) reviews this evidence in detail, together with the two properties of the underlying measure that motivate our approach.

This powerful body of work rests almost entirely on analyses of **functional connectivity**, defined as the correlation or statistical dependence between regional time series. As an undirected quantity, functional connectivity cannot discriminate between competing causal architectures. For instance, an elevated thalamus–somatosensory correlation in schizophrenia is equally compatible with thalamic drive of cortex, cortical drive of thalamus, or modulation by a common source. Nevertheless, the interpretive framework commonly applied to such findings, invoking thalamic overdrive or prefrontal regulatory failure, implicitly assumes directed influences. This directionality is, however, more often inferred in practice from anatomical priors and theoretical commitments, rather than directly from the data. Furthermore, correlation can appear in regions that are not directly connected.

**Effective connectivity** methods aim to supply that missing direction, but in fMRI they confront a fundamental obstacle inherent to the measurement method itself. The **blood-oxygenation-level-dependent** (BOLD) signal is sampled in the order of seconds, which is approximately one order of magnitude slower than the neuronal interactions it indexes [25, 31]. A causal graph estimated at this measured timescale is therefore a low-pass filtered image of a faster underlying process. Moreover, this hemodynamic convolution is not a benign smoothing: it can hide true edges, fabricate spurious ones, and most consequentially for directional claims, reverse the apparent orientation of an edge or even entire circuit [16, 36, 22]. Methods that commit to the measured timescale, treating one BOLD sample as one causal step, implicitly inherit these distortions. Directional estimates are therefore particularly sensitive to model misspecification at the measured BOLD timescale.

A line of work that is agnostic to signal undersampling rate addresses this problem [16, 36, 22, 1]. Not assuming that the measured graph equals the neural one, these methods treat the sampling rate as unknown and recover the set of latent-timescale graphs consistent with the observed data, explicitly modeling how undersampling maps a fast causal process onto a slow measurement. We build on this framework with RnR, an undersampling-aware causal discovery method [2] that refines the graph returned by a first-order estimator by accounting for undersampling. RnR returns an **equivalence class** of plausible directed graphs, each annotated with the sampling rate that would produce it, rather than a single graph that can signal unjustified certainty.

Here we bring this tool to the schizophrenia connectome. Our aim is to investigate whether estimating directed causal graphs using the RnR approach on a large schizophrenia dataset reproduces the established functional connectivity picture, expresses it in directed form, and clarifies where directionality can and cannot be trusted. We applied RnR to resting-state fMRI from the multi-site Function Biomedical Informatics Research Network (FBIRN) cohort [24], the same source data and independent-component template family [17] that underlie the canonical dynamic-connectivity reference study [15]. Specifically, we (i) recover a directed patient-versus-control connectome and show that it reproduces the field’s most replicated finding— the sensory/thalamic hyperconnectivity hub—in oriented form; (ii) benchmark this recovery against five single-timescale estimators; (iii) classify each directed edge by the stability of its orientation across the inferred undersampling rate, distinguishing directions that are timescale-robust from those that are timescale-contingent, and find every reported edge to be robust at this resolution; and (iv) show through simulation how spatial resolution impacts the perceived undersampling rate, explaining why single-timescale methods have succeeded and when they will fail. Where our directed findings depart from the correlational literature, they do so almost exclusively on couplings the literature itself reports as unstable, or on orientations that an undirected measure cannot specify. Thus, modeling temporal undersampling changes what can be recovered from fMRI, and specifically what can be recovered about direction. The synthetic experiments establish that claim where ground truth exists, the FBIRN analysis tests it where ground truth does not, and the resolution simulation explains why the field has been able to proceed without it so far.

### Interpretational scope of the directed claims

Because the vocabulary of causal discovery is stronger than the vocabulary of correlation, we state at the outset what a directed edge in this paper does and does not assert. An edge *i* → *j* is a statement within the model class defined in Section 3.1: in the recovered causal-timescale graph, component *i* at one timestep is a cause of component *j* at the next, in the standard interventionist sense [44] that there exist conditions in which manipulation of *i* would alter the distribution of *j*. Three qualifications attach to every such statement. First, the claim is conditional on the modelling assumptions, namely first-order dynamics at the latent scale, stationarity within scan, and causal sufficiency at the component level up to the hidden common causes that undersampling renders as bidirected edges. Second, the units are spatially independent components, each a distributed weighted map rather than an anatomical region, so an edge is a relation between components and not evidence of a monosynaptic projection between the structures at their peaks. And third, no universal interventional claim is intended: we do not assert that manipulating *i* will always change *j*, only that there exist conditions in which it would. Given these assumptions, some causal directions may be underdetermined by the data, so we label each orientation by its stability across the inferred sampling rates (Section 3.4) to distinguish orientations required by measured data versus those left open.

## 2 Related Work: Replication Gaps and Connectivity

We begin by reviewing robust connectivity findings from two decades of fMRI research, then specify the two properties of the underlying measure that our approach addresses: undirectedness and single-timescale sampling.

### 2.1 The thalamocortical signature

The most reproducible resting-state finding in schizophrenia is a thalamocortical dysconnectivity motif with an unusually consistent form: reduced thalamo-prefrontal coupling together with increased thalamo-sensory and thalamo-sensorimotor coupling. **(author?)** [45] first characterized this pattern by parcellating the cortex and mapping its connectivity to the thalamus, reporting reduced prefrontal–thalamic and increased motor/somatosensory–thalamic connectivity. **(author?)** [4] replicated it with anatomically defined thalamic seeds in ninety patients, adding thalamic under-connectivity with prefrontal–striatal–cerebellar regions and interpreting the combined picture in terms of disturbed sensory gating and top-down control. A multi-site brain-wide association study of 415 patients and 405 controls confirms this pattern, identifying the thalamus as the principal aberrant hub, with increased thalamus-somatosensory connectivity the most significant group difference and thalamo-frontal connectivity weakened [13]. Independent component analyses (ICA) of multi-site resting-state data likewise recover thalamic hyperconnectivity with auditory, motor, and visual networks alongside reduced connectivity among sensory networks [15]. The robustness of this finding across seed-based, ICA, and dynamic methods, and across chronic, first-episode, and clinical-high-risk samples, has led several authors to treat thalamocortical dysconnectivity as a candidate core neurobiological signature of the illness.

### 2.2 Cortical networks: default-mode, salience, and segregation

Two themes recur at the level of canonical cortical networks. First, within-network hypoconnectivity is common in the default-mode and self-referential systems, frequently implicating anteromedial hubs (anterior cingulate, medial prefrontal) and posteromedial hubs (posterior cingulate, precuneus). Second, between-network coupling is disturbed, particularly in the interactions of the salience/ventral-attention system with the default-mode and executive systems, along with a broad reduction in the normal segregation between task-positive and task-negative networks. The salience network is central to this account. On the influential triple-network view, the right fronto-insular cortex acts as a switch that toggles between the default-mode and central-executive networks [42], and failure of this switching has been proposed as a mechanism for the disrupted network dynamics of schizophrenia [29]. Cerebellar involvement recurs as well, consistent with the classic “cognitive dysmetria” account of a disrupted cortico-subcortical-cerebellar circuit [3].

### 2.3 A recurring interpretive tension: hyper-versus hypoconnectivity

A persistent complication is that “hyperconnectivity” and “hypoconnectivity” are not fixed labels. Their sign depends on illness stage (early-course cohorts more often show prefrontal hyperconnectivity, chronic cohorts hypoconnectivity), methodology (seed-based versus ICA, static versus dynamic), medication exposure, and preprocessing choices such as motion handling and global-signal treatment [15, 34]. The same pair of regions can be reported as hyperconnected in one design and hypoconnected in another. This sign instability arises from an undirected measure applied across heterogeneous samples and pipelines, and it affects how strongly any single reported difference should be weighted.

### 2.4 Two intrinsic limits of functional connectivity: direction and timescale

Two properties of functional connectivity underlie both its successes and its ambiguities. The first is that it is undirected. A reported group difference is a directionless difference in the strength of statistical dependence between two regions. The directional interpretations the field routinely attaches—thalamus drives sensory cortex, and prefrontal cortex fails to exert control—are imported from anatomy and theory, not inferred from the data. The second is that functional connectivity is estimated at a single, slow timescale. BOLD is sampled far more slowly than neural dynamics, so any dependency structure derived from it reflects a temporally coarse view of the underlying process [25]. When direction is estimated at all, this coarse temporal sampling can reverse it [16]. These two limits are exactly what a causal discovery method accounting for undersampling sets out to address, and they guide our empirical approach. The central question of this work is not whether the schizophrenia connectome is disrupted, as that is well established, but what it looks like when rendered as a directed graph that treats slow sampling as a first-class part of the model.

## 3 Methods

Our goal is to infer the true causal graph *G*^1^ given only a measurement graph ℋ obtained under temporal undersampling, where the rate of undersampling *u* is not known in advance. Throughout this paper, we use “measurement” timescale to refer to slower/measurable data, and “causal” timescale to refer to faster underlying system respectively. Figure 1 gives an overview of the full pipeline, from 4D fMRI volumes and first-order graph estimation through to the undersampling-aware equivalence class of directed graphs we report.

**Figure 1.**
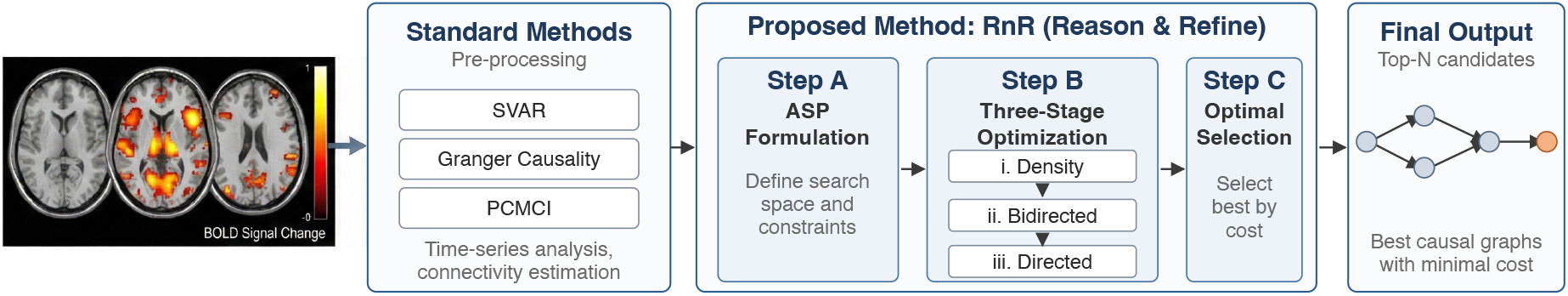
Overview of our pipeline. Starting from 4D fMRI volumes, classical methods extract a single “measured” graph from BOLD time series (e.g., using GC, SVAR, or PCMCI). These approaches typically ignore the effects of temporal undersampling. Our method extends beyond this step by accounting for the ambiguity introduced by slow sampling. Instead of selecting a single best-fit graph, we recover an equivalence class of plausible causal graphs that are consistent with both the observed data and the structural distortions caused by undersampling. Finally we present top *N* graphs based on optimization cost.

A directed dynamic causal model extends standard causal models [33] by adding time: *n* random variables **V** = {*V*_1_, …, *V*_*n*_} appear at both the current timestep *t* (**V**^*t*^) and previous timesteps (**V**^*t*−*k*^). The full dynamic graph **G** is defined over 2**V**, and the only permissible edges are 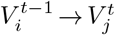 (including *i* = *j*). We assume first-order Markov structure: **V**^*t*^ *⫫* **V**^*t*−*k*^ | **V**^*t*−1^ for all *k >* 1.

Suppose the causal process unfolds over the timepoint sequence {*t*^0^, *t*^1^, *t*^2^, …} but observations are recorded only every *u*-th point, i.e., at {*t*^0^, *t*^*u*^, *t*^2*u*^, …}; we then say the data are undersampled at rate *u*. Traditionally, BOLD fMRI samples the brain at a repetition time (TR) which is in the order of seconds. That is one to two orders of magnitude slower than the neural interactions it indexes [31, 25]. We model the latent neural process as a first-order dynamic causal system. Let *G*^1^ be its directed graph at the causal timescale, with binary adjacency **A** ∈ {0, 1}^*n×n*^. A graph ℋ, read at the slow measured scale is a temporally subsampled view of this faster process, and subsampling can hide edges, add edges, and flip edge direction [16, 36, 22].

A compressed graph *G* represents [16] a dynamic graph **G** over the *n* variables alone, with temporal information encoded implicitly in the edges: a directed edge (*i* → *j*) marks a directed walk of length *u*, and a bidirected edge (*i* ↔ *j*) marks two nodes reached from a shared ancestor *w* by directed walks of equal length *ℓ < u*:

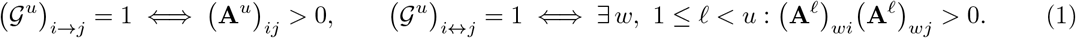

Recovery reverses this operator while being agnostic to the rate. Given a measured graph ℋ, Rate-Agnostic Structure Learning (RASL) returns all causal-timescale graphs with rates that could produce ℋ [36]:

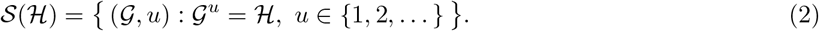

This inverse operation is NP-complete and typically yields multiple (graph, rate) pairs, so RASL returns the full equivalence class, not one graph [22]. Solver-based RASL (sRASL) [1] writes the inversion as an Answer Set Programming (ASP) problem. ASP is a declarative form: one states logical constraints, and a solver lists every graph that meets them. sRASL exploits a key structural fact to dramatically speed up recovery. Specifically, the node membership of a strongly connected component (SCC)—a maximal set of nodes such that there is a directed path from every node to every other—is almost always stable under undersampling. This stability requires only that the greatest common divisor of the set of simple loop lengths in the SCC is one; this is a weak assumption that holds, e.g., if there is a single auto-correlated variable in the SCC. Moreover, the edges between SCCs necessarily form a directed acyclic graph (DAG). These properties jointly impose significant additional constraints that sRASL can exploit to prune much of the search up front. Search can also be simplified since there is a rate past which more undersampling adds no new graphs [36]; for fMRI, a loose bound is near *u*=20, well above the rates we find.

### 3.1 Undersampling-aware causal discovery (RnR)

In this section, we present Real-world noisy RASL (RnR), our enhanced structure learning framework that builds upon the sRASL [1] framework through five key innovations tailored for noisy, undersampled fMRI data. While each individual piece might appear relatively simple, they collectively have a large positive impact on causal structure learning from real-world data. The strength is in how these pieces fit together, each one shaped for a difficult, pressing scientific problem:

1. To address the possibility that the least-cost graph is not always the closest graph to the underlying causal scale ground truth, RnR returns every candidate whose cost stays within a relative tolerance of the best: *C*(*G*) ≤ Δ with Δ = 1.9 *c*_min_, where *c*_min_ (possibly more than one candidate with cost= *c*_min_) is the lowest cost found. The tolerance is relative, so it scales with graph size. This set is the working stand-in for *S*(ℋ).
2. Realistic brain networks are neither near-empty nor near-full. Their edge density sits in a moderate band, roughly 10 to 30 percent of possible edges [41, 30, 6]. RnR sets a target density in this band and imposes a cost on candidates based on the distance of their density from that target, so solutions stay biologically plausible.
3. The cost is split into three components, *C*(*G*) = *C*_*o*_ + *C*_*b*_ + *C*_*d*_: a density term, a bidirected-structure term, and a directed-orientation term. RnR minimizes them in that fixed order by first minimizing density error, then errors on bidirected links, followed by edge direction. The optimization order thus runs from low-information to high-information. Each stage cuts the candidate set by orders of magnitude before the hard directional choices are made.
4. RnR weights each edge by its evidence, as not every proposed edge or absence has equal support. The solver pays a large cost to drop a strongly supported link and a small cost to drop a weak one, as well as a high default cost to add any edge the first-order method did not propose. Here, we derive the weights directly from the underlying causal structure estimating methods that we use (e.g., PCMCI, Granger causality, MVAR, etc.), though weights can also encode prior anatomical knowledge. Recovery is sensitive to evidential weights, not uniform across possible edges.
5. RnR can be used as a meta solver on top of existing methods, to add undersampling-aware treatment to the task at hand. Undersampling turns a hidden common cause into a bidirected edge (Eq. (1)). A first-order estimator with no bidirected primitive cannot draw that edge, so it splits the same dependence into a mutual 2-cycle, *i* → *j* and *j* → *i*. That two-cycle is the footprint the confounder leaves in the output for the first-order estimator, here the PCMCI method. So for each mutual cycle in ℋ, RnR offers the solver a bidirected edge alongside the two directed edges, with a low penalty on the directed pair. It lets the cost pick the pre-image: bidirected only, one direction, a true 2-cycle, or all three. This folds undersampling awareness into the upstream output.

Therefore, RnR’s workflow as illustrated in Figure 1 becomes: per participant, we first estimate ℋ with PCMCI [38] (ParCorr test, lag *τ*_max_=1, *α*_PC_=0.01). We convert the lagged PCMCI output to a compressed causal graph and solve Eq. (2) with the `gunfolds` [35] implementation of RnR under the domain-constrained SCC encoding. RnR returns a set of (*G, u*) pairs per participant. We pool the *k*=3 lowest-cost members. Realized rates fall in *u* ∈ {2, 3, 4, 5}, with about 78% of solutions at *u*=2. The low rate fits the voxel-timescale argument: a voxel holds 8 to 10 layers of neurons, so its causal timescale is plausibly about 1000 ms, which corresponds to *u*=2. We expand this idea later in Section Section 5. A domain-specific description of the ASP predicates, the exact sRASL encoding, RnR’s weighted relaxation, and the conversion of `clingo` answer sets back into (*G, u*) pairs is provided in Section 7.

### 3.2 Group-difference edge test

A principal application of causal discovery in neuroscience is the characterization of differences between distinct populations. Clinical investigations of many neuropsychiatric disorders rely on the assumption that diagnostic categories and typical control samples each have within-group homogeneity. Studying the brain connectivity of groups is crucial for several tasks, such as detecting diseases and disorders, designing effective therapeutic interventions, and mapping brain networks. We adopt this view here and test for group-level differences edge by edge.

For each ordered pair of nodes (*i, j*) we tested whether the presence of the directed edge *i* → *j* differs between groups. For RnR we pooled each participant’s *k*=3 lowest-cost solutions (its most probable members of the undersampling equivalence class), giving pooled group sizes *n*_0_ = *k* |HC| = 453 and *n*_1_ = *k* |SZ| = 480; each baseline contributes its single graph, so *n*_0_=151, *n*_1_=160. Let *a*_*ij*_ and *b*_*ij*_ be the number of pooled HC and SZ graphs containing *i* → *j*. We formed the 2 × 2 table 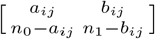 and computed an exact (Fisher) *p*-value *p*_*ij*_, with signed effect size

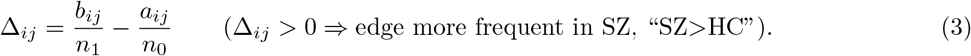

Across the *m* = *N* (*N* − 1) off-diagonal edges (*m*=90 at *N* =10) we controlled the **false discovery rate** by Benjamini–Hochberg [**(year?)**]: sorting *p*_(1)_ ≤ · · · ≤ *p*_(*m*)_, we declared significant all edges up to the largest rank *k*^⋆^ with 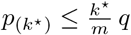 at *q*=0.05. We report both the FDR-significant set and a “suggestive” set (*p*_*ij*_ *<* 0.05, uncorrected). We use the exact Fisher test for every cell.

### 3.3 Literature-anchored scoring

To distinguish “more edges” from “more credible edges,” we scored each significant edge set *E* against a curated map L of undirected SZ-vs-HC connectivity pairs, each tagged with an expected coupling sign (hyper-or hypoconnectivity, i.e. SZ*>*HC or SZ*<*HC) drawn from the FNC literature (see Section 5). For the subset of pairs testable in a given node set, ℒ_*T*_, we defined

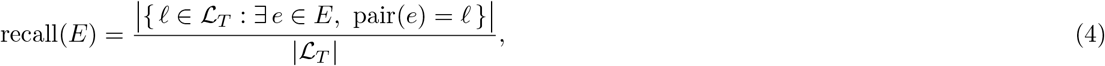

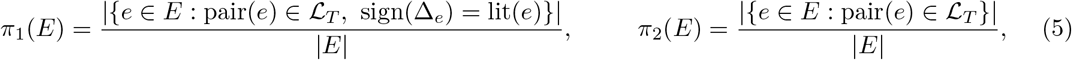

where recall measures coverage of the known pairs. sign(Δ_e_) is the effect sign of edge *e* (hyper-vs. hypoconnectivity, i.e. SZ*>*HC vs. SZ*<*HC) and lit(·) the literature’s expected coupling sign for that pair; both are magnitude signs, not causal orientations. Thus, *π*_1_ (sign-concordant precision) requires the pair to be documented and its hyper-/hypoconnectivity sign to match the literature, whereas *π*_2_ drops the sign match and keeps any edge for a documented pair. We relax the sign requirement because the literature’s hyper-/hypo-connectivity judgment is a property of the whole undirected pair (its net coupling magnitude) and cannot be attributed to a single directed limb, so a sign mismatch on one directed edge is not a clean contradiction. We additionally define an undersampling-aware precision *π*_3_ in Section 3.4.

### 3.4 Undersampling robustness of edge direction

RnR enables such questions as whether an edge’s orientation is an artifact of the assumed timescale. Pooling all solutions across all participants and partitioning by their inferred rate *u*, we counted, for each pair (*i, j*) at each *u*, the solutions in which exactly one orientation is present (“directionally resolved”), and computed

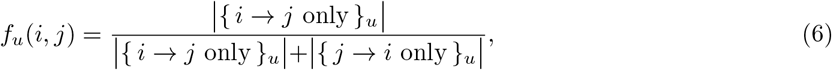

with dominant orientation *i* → *j* if 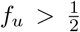. A rate level “votes” only if it has ≥ 30 resolved solutions. The consistency of a pair is the fraction of voting levels whose dominant orientation matches the pooled-dominant orientation. We label a pair **undersampling-robust** if consistency = 1 (orientation invariant across all voting *u*), with ≥ 2 voting levels and directed presence ≥ 0.10; and **undersampling-fragile** if the dominant orientation flips across voting levels (consistency *<* 1) with decisive margins 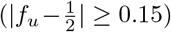. Pairs meeting neither definition, too few voting levels, directed presence below 0.10, or a dominant orientation that changes only within the indecisive band 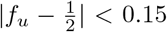, are left unclassified. Finally, the undersampling-aware precision credits any edge except a <u>hard contradiction</u>—an edge that is sign-discordant with and undersampling-robust (so a direction flip cannot explain the disagreement):

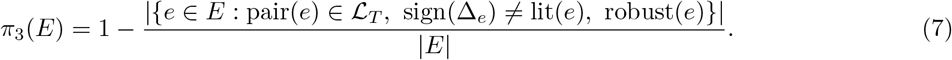

### 3.5 Synthetic validation with known ground truth

Because the true differential connectome is unobservable in fMRI, we validated the discovery procedure on synthetic data where the group difference is known by construction. We generated repeated random pairs of six-node graph families (*G*^HC^, *G*^SZ^) that share a strongly connected backbone and differ in a fixed set *D* of directed edges (edges gained or lost in “SZ”). For each synthetic participant we drew a stable vector-autoregressive process on the group graph and sampled it every *u* steps, so the observable is a realization of the order-*u* footprint *G*^u^ (Eq. 1). We then applied the same RnR pipeline used on the real data (PCMCI → RnR) together with a panel of five single-graph baselines fixed at the measured timescale (*u*=1), and scored each method’s recovered edges against the known *D* by recall, precision, and *F*_1_ as a function of pooling depth *k*. To keep the comparison fair, each baseline was inflated to the same *k* by pooling *k* hyperparameter-jittered fits, so that any dependence on *k* reflects the content of the multiplicity rather than a larger sample; the unit of analysis is the participant (one soft edge-probability vector per subject), not the individual solution.

The design targets a limitation intrinsic to undersampling rather than any particular estimator. A single-graph method sees only the order-*u* footprint G^u^ (Eq. 1), so it is blind to any edge differences that appear the same across the two groups; writing *M*_u_ ⊆ *D* for these <u>masked</u> edges, the population recall of any *u*=1 method cannot exceed 1 − |*M*_u_ | */* | *D* |. Because the footprint of a strongly connected, aperiodic graph saturates toward the complete digraph as *u* grows [21], masking eventually claims every edge difference and maximum recall falls to zero. In contrast, RnR reasons over the equivalence class () of latent graphs and rates (Eq. 2), so can theoretically maintain high recall.

## 4 Results

We evaluate RnR in three stages, from full ground-truth control toward real data. Section 4.1 works with synthetic time series, where the generating graph and the undersampling rate are both known: we benchmark against an established fMRI simulation benchmark and against more general VAR-driven graphs, quantify how performance degrades as the sampling interval lengthens under a balloon-model BOLD forward pass. Section 4.2 turns to real fMRI, contrasting RnR with Granger Causality Mapping on a well-studied six-component network to show what an undersampling-aware view adds to a purely temporal one. Section 4.3 applies the method to resting-state fMRI from 311 FBIRN participants, where the true connectome is unobservable and the claims must instead be supported indirectly: by the amount of differential structure recovered relative to single-timescale baselines, its agreement with the connectivity literature and with other methods, and its stability across inferred sampling rates.

### 4.1 Synthetic data

Our initial evaluation of RnR used the synthetic benchmark introduced by **(author?)** [39]. We show that the approaches examined in that work, including methods designed specifically for this benchmark, fail to address the distorting impacts of undersampling.

We benchmarked RnR against five single-graph estimators that commit to the measured timescale (*u*=1): PCMCI [38]; FASK [39]; GIMME [19]; a multivariate autoregressive (MVAR) model [20]; and conditional multivariate Granger causality (MVGC; 7, 8). Each baseline outputs one binary directed graph per participant, in the identical adjacency convention used downstream. Figure 2 summarizes these comparisons across all networks. The green bars report the performance we obtained by running the official implementation of each method ourselves and scoring the output against the true generating graph. The orange bars, labeled GT^2^, score those same outputs against the ground truth undersampled by a factor of two; their consistently higher values indicate that, precisely because these methods do not model undersampling, their estimates align more closely with the undersampled graph than with the true one. The purple bars (meta-RnR) show the result of applying RnR as a meta-solver on top of each method’s output, while the pink bars (RnR) correspond to our full pipeline of RnR applied to PCMCI, the base structure-learning method we found to pair best with our approach.

**Figure 2.**
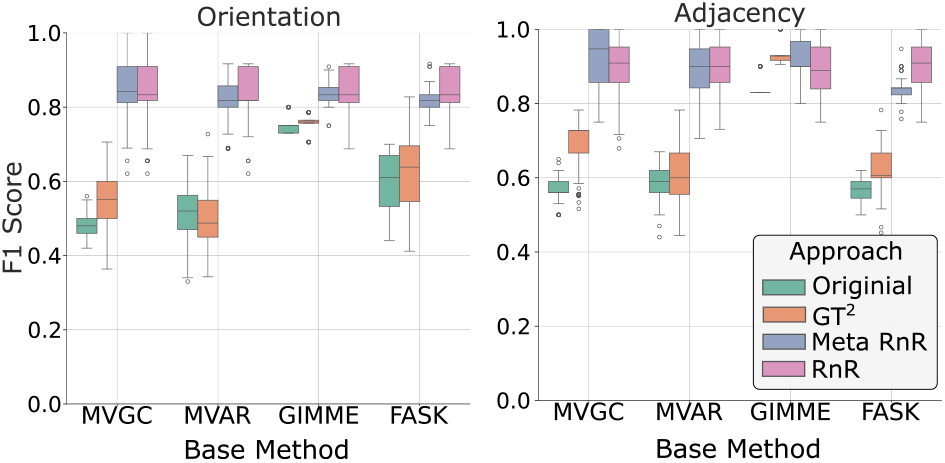
Comparison of performance on Sanchez-Romero’s data with and without the RnR meta-solver. The original methods score higher against the ground truth undersampled by two (GT^2^, orange) than against the true graph (green), confirming that they ignore undersampling. Layering RnR over these methods yields higher accuracy by correcting for undersampling effects (purple), and combining RnR with PCMCI provides further improvement (pink).

Although Sanchez-Romero’s data-generation approach has been used in previous evaluations, it is limited in both scope and complexity: it relies on small, simple graphs that lack genuine loops and are generated using a specific simulation procedure. To evaluate performance on a broader graph family for more general causal inference, we ran experiments with VAR models on randomly generated ring graphs (Figure 3). The top two panels report simulations on a random ring graph that has additionally been passed through a VAR simulation, while the bottom two panels use Sanchez-Romero’s simple network and seed graph with a VAR simulation applied on top. Our method performs better on both types of data; the competing methods, by contrast, suffer a marked drop in F1 across both orientation and adjacency once they are applied to random, general graphs. This highlights the importance of testing causal inference methods on more realistic and complex data.

**Figure 3.**
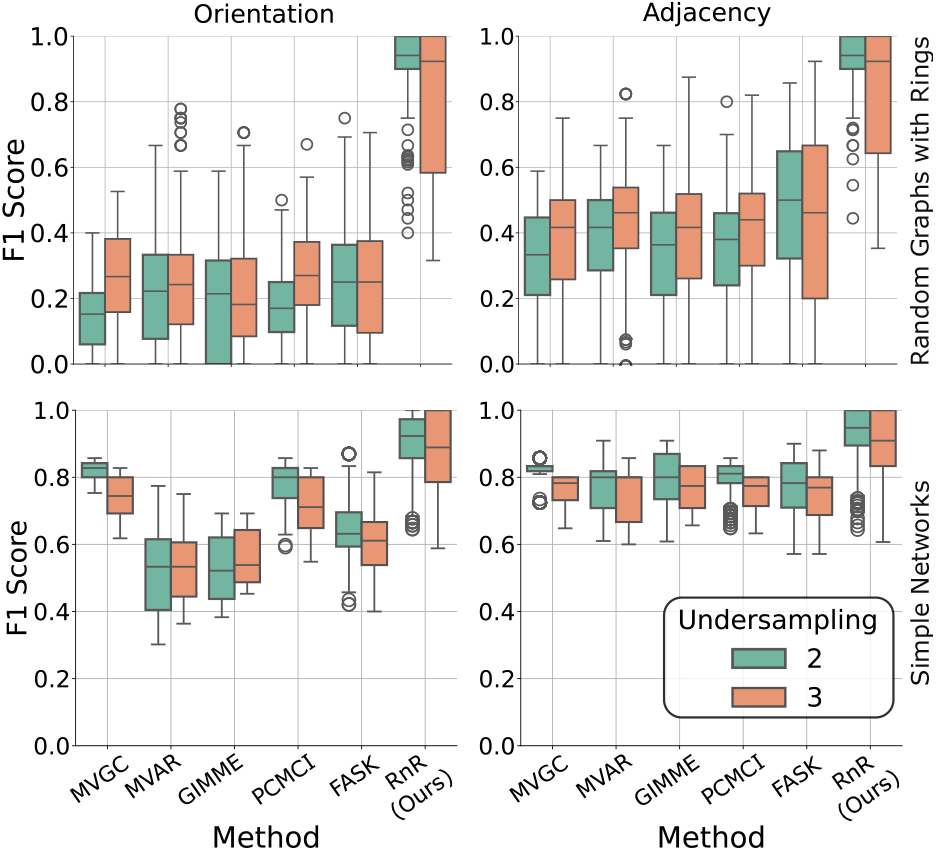
Comparison of Sanchez-Romero’s simple data-generation scheme with larger, more generalized VAR-generated graphs. The top two panels show a random ring graph and the bottom two Sanchez-Romero’s seed network, each processed through a VAR simulation. Because Sanchez-Romero’s data lacks real loops and is narrow in scope, it may limit the generalizability of causal inference results: the baseline methods lose substantial F1 in both adjacency and orientation on the random general graphs, whereas RnR remains the strongest performer across both data types.

We also studied the effect of undersampling on time series generated from a VAR model and then passed through a balloon model to emulate the BOLD response in fMRI data [10]. Because the BOLD signal is inherently smooth, undersampling at one-second intervals typically incurs minimal information loss, whereas undersampling at larger intervals (e.g., two or three seconds) introduces substantial distortion. Figure 4 shows how different undersampling rates affect the preservation of temporal information in BOLD data: for every method that ignores undersampling, both graph accuracy and F1 degrade as the undersampling rate grows, whereas our method remains comparatively stable and robust. This underscores that aggressive undersampling can obscure meaningful connectivity patterns. Across these experiments (Figures 3, 4), RnR improves F1 by 64% on average over the baselines.

**Figure 4.**
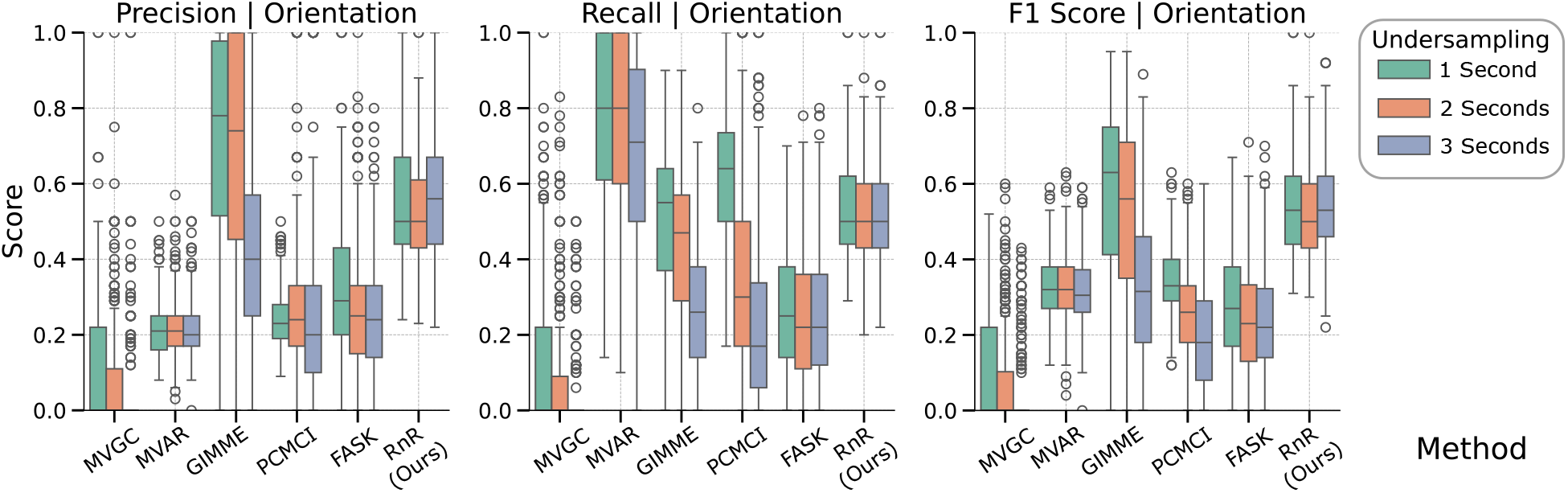
Impact of different undersampling rates (1s, 2s, 3s) on BOLD signal preservation in fMRI data simulated with the balloon model. Minimal error is observed with 1-second undersampling, whereas larger intervals degrade accuracy in all methods that don’t account for the undersampling effect. Our method RnR accounts for this effect and does not suffer loss from undersampling.

Finally, we ran a synthetic experiment to assess performance not on recovery of a single graph, but rather on the <u>difference</u> between two of them. In other words, we want to know if an undersampling-aware treatment of time series data can help us better differentiate between two groups of underlying models. Because the true differential connectome is unobservable in fMRI, we first confirmed on synthetic data with a known group difference (Fig. 5) that RnR’s recovered difference edges are genuine rather than artifacts of pooling multiple solutions. We generated 12 randomly generated six-node graph pairs separated by a known set of edges, a controlled stand-in for the healthy-control versus schizophrenia (HC/SZ) contrast we analyze later. Across these pairs, RnR’s recovered difference was the most accurate at every pooling depth *k*: its precision and F1 against the ground-truth difference were 2–3 × those of every single-graph baseline, and its recall approached the most liberal baseline (PCMCI) by *k* ≈ 8, a level PCMCI reaches only by flagging far more edges at a third of the precision. The baselines were flat in *k* (inflating each to the same number of pooled solutions did not improve it), whereas RnR moved with *k*, confirming that the gain derives from the diversity of the undersampling equivalence class rather than sample-size inflation.

**Figure 5.**
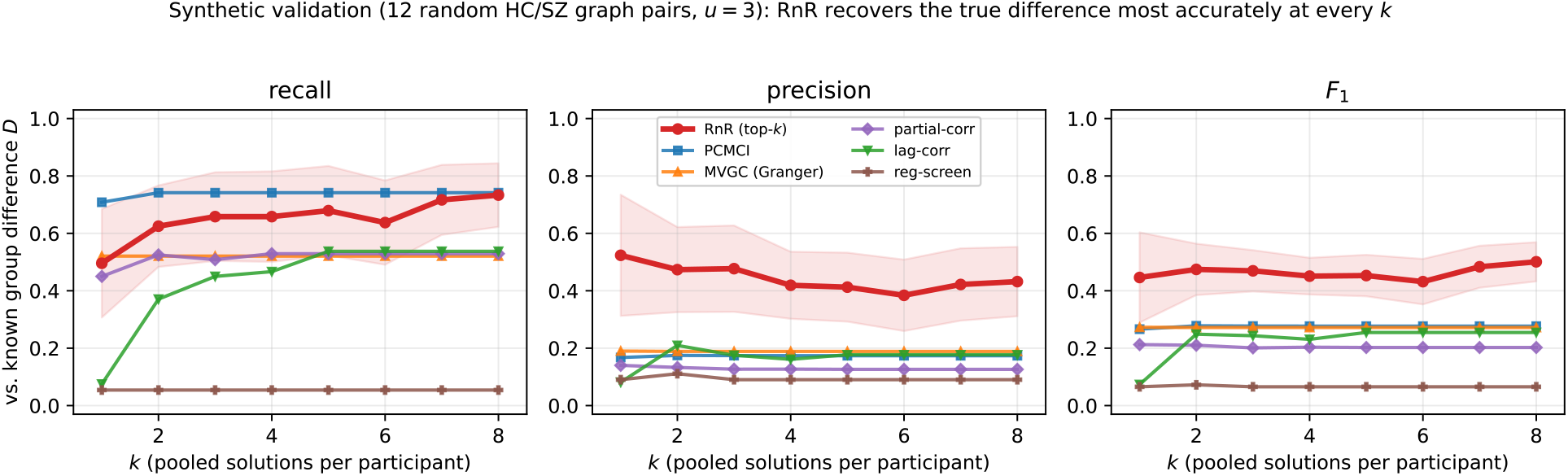
Synthetic validation on a known group difference. Recall, precision and *F*_1_ of the recovered difference edges versus pooling depth *k*, over many random six-node HC/SZ graph pairs observed at *u*=3 (RnR in red; five single-graph baselines each inflated to the same *k*). RnR attains the highest precision and *F*_1_ at every *k* (2–3 × the baselines); its recall approaches the most liberal baseline by *k* ≈ 8. The baselines are flat in *k* (inflation adds nothing), whereas RnR improves with the diversity of the equivalence class.

Together with the rate-recovery experiments above, these simulations provide a ground-truth evaluation that is unavailable in the cohort analysis: where the sampling rate departs from the causal timescale, single-timescale methods recover systematically biased structure, whereas RnR’s pooled, undersampling-aware estimate recovered more of the simulated differential structure as additional candidates were included. The more demanding test is whether this advantage survives the move to real data, where the generating connectome is unobservable and the true undersampling rate unknown. There, the value of undersampling-aware discovery cannot be read off a ground-truth graph and must be established indirectly, through reproducibility of known effects, internal consistency across inferred timescales, and agreement with the connectivity literature. In the next sub-section, we apply RnR to resting-state fMRI from a large, multi-site schizophrenia cohort.

### 4.2 Comparative Analysis of Causal Discovery Methods on fMRI Data

To validate our causal discovery framework and demonstrate its ability to extract novel information from neural imaging data, we conducted a comparative analysis using Granger Causality Mapping (GCM) [37] and our RnR. We utilized the same six ICA-derived brain regions as [42], spanning the Central Executive Network (CEN), Salience Network (SN), and Default Mode Network (DMN). By paying close attention to causal rates and sampling distinct causal properties, our results reveal that different methods can uncover complementary layers of network organization.

We first applied GCM to validate our pipeline against established findings. Our in-house implementation (Figure 6 left) showed substantial agreement with [42]’s results. Specifically, we confirmed the central role of the right fronto-insular cortex (rFIC) in mediating network interactions. As shown in Figure 6 (left) and Table 1, the strongest connection was the coupling between rFIC and the right dorsolateral prefrontal cortex (rDLPFC), replicating the key salience-executive link emphasized in the original study. Furthermore, we replicated the finding that the Salience Network exerts influence over the DMN, with robust directed connections from rFIC and ACC to DMN regions (ventromedial prefrontal cortex (VMPFC) and posterior cingulate cortex (PCC)). These agreements validate the fidelity of our implementation and the quality of the underlying data processing.

**Table 1.** Top-5 causal connections identified by GCM (temporal) and RnR (structural).

| GCM (Temporal Causality) |  |  | RnR (Structural Causality) |  |  |
| --- | --- | --- | --- | --- | --- |
| Connection | Freq. | Interaction Type | Connection | Freq. | Interaction Type |
| rFIC ↔ rDLPFC | 58.1% | Saliency ↔ CEN | VMPFC → rFIC | 89.7% | DMN → Saliency |
| VMPFC → PCC | 39.4% | DMN (internal) | rDLPFC → ACC | 67.3% | CEN → Saliency |
| PCC → VMPFC | 39.0% | DMN (internal) | PCC → rFIC | 64.1% | DMN → Saliency |
| VMPFC → rDLPFC | 38.4% | DMN → CEN | VMPFC → PCC | 62.6% | DMN (internal) |
| rDLPFC → VMPFC | 37.4% | CEN → DMN | VMPFC → rPPC | 59.1% | DMN → CEN |

**Figure 6.**
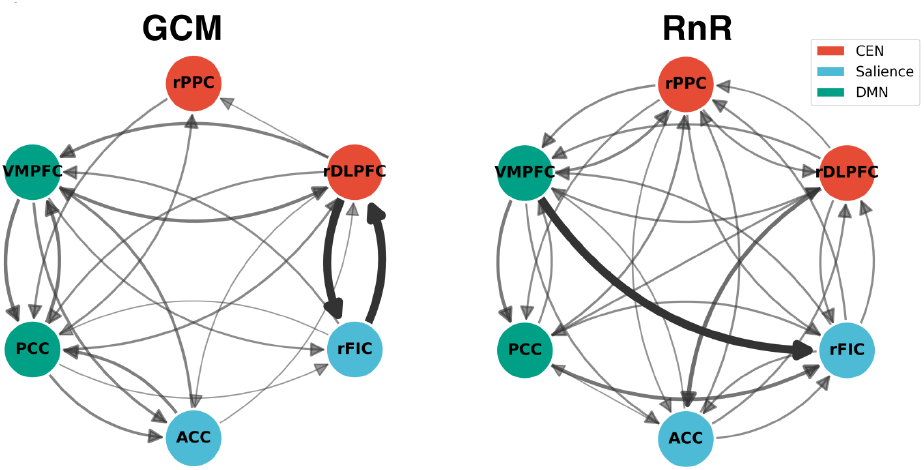
Comparative analysis of GCM and RnR. **Left:** GCM network showing strong bidirectional rFIC– rDLPFC coupling (thick black edges), replicating [42]’s salience–executive findings. **Right:** RnR network revealing a dominant unidirectional pathway from the DMN, suggesting a structural hierarchy absent in the bidirectional GCM results.

Applying our RnR method to the exact same ICA components yielded a different but related set of answers, highlighting the method’s sensitivity to structural rather than purely temporal causal signals. While GCM emphasized bidirectional coupling (rFIC ↔ rDLPFC), RnR identified a hierarchical structure driven by the Default Mode Network (Figure 6 right). The most significant finding was a robust unidirectional connection from VMPFC to rFIC (Figure 6 right, Table 1), which was present in nearly 90% of solutions—a far stronger signal than any individual connection found by GCM.

Comparing the two approaches reveals important convergences and one critical divergence. Both methods agree on the high level of interconnectivity between the three networks and the central importance of the Salience Network (rFIC/ACC) in bridging distinct functional systems. However, the methods disagree on the primary driver of the system. GCM identifies the reciprocal temporal dynamics between rFIC and rDLPFC as the dominant feature, likely reflecting the active “switching” mechanism. In contrast, RnR identifies the structural precedence of VMPFC over rFIC, suggesting that the DMN’s state may structurally constrain or gate the activity of the Salience Network. Modelling the undersampling effect thus reveals structure that the temporal view misses.

### 4.3 Application to Schizophrenia fMRI

#### 4.3.1 Participants and functional network nodes

We analysed resting-state fMRI from the multi-site Function Biomedical Informatics Research Network (FBIRN) phase-III schizophrenia study [24]: 311 participants, 151 healthy controls (HC) and 160 individuals with schizophrenia (SZ). Network nodes were defined as spatially independent components from the <u>NeuroMark_fMRI_1.0</u> group-ICA template [17, 11], which assigns each component to one of seven functional domains (subcortical SC, auditory AU, sensorimotor SM, visual VI, cognitive control CC, default-mode DM, cerebellar CB). The same source cohort (FBIRN phase-III) and group-ICA lineage underlie the canonical static/dynamic functional-network-connectivity (FNC) reference study of **(author?)** [15], making our component-level results comparable at the component/domain level to that work.

Components were selected according to three criteria: (i) coverage of all seven NeuroMark functional domains; (ii) prioritization of those regions most robustly implicated in the schizophrenia functional-connectivity literature: the thalamus and striatum, primary visual, auditory and sensorimotor cortex, anteromedial and posteromedial default-mode hubs, the salience-network insula, and the cerebellum [45, 15, 13]; and (iii) a node count within the range where undersampling-aware causal discovery remains computationally tractable, since exact recovery scales exponentially with the number of nodes. These criteria yielded a 10-node set (*N* =10) that spans all seven functional domains.

#### 4.3.2 RnR recovers the richest differential connectivity

Under one common test on the same 311 participants, RnR recovered far more FDR-significant HC-vs-SZ directed edges than any comparison method (Fig. 7): 14 edges, versus ≤ 3 for every other method (PCMCI 2; MVAR 3; FASK 2; GIMME 0; MVGC 0). The baselines are not insensitive: at the suggestive threshold, every method detected some group structure (RnR 26 edges; others 6–10), so the methods differed in the number of edges crossing the selected thresholds. The single-graph methods produce one graph per participant (*u*=1); RnR’s larger yield is a direct consequence of integrating over a full undersampling equivalence class.

**Figure 7.**
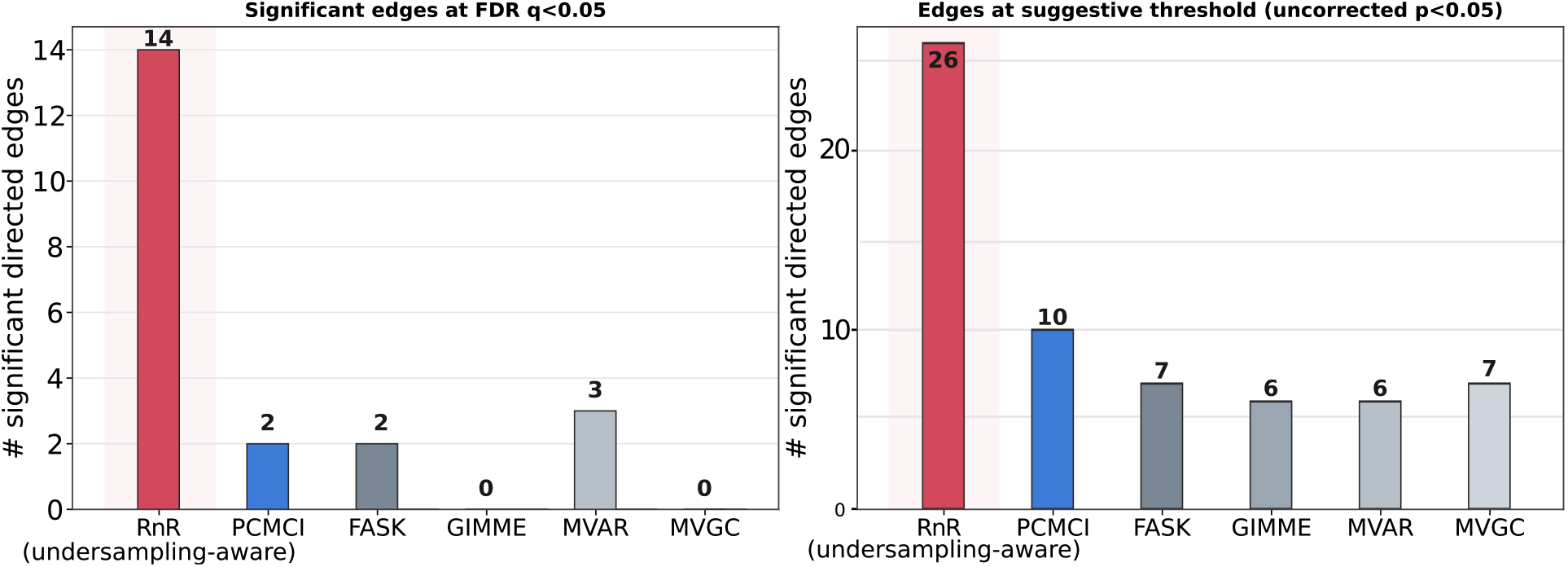
RnR recovers the most differential connectivity. Number of HC-vs-SZ directed edges per method, at FDR *q<*0.05 (left) and the suggestive threshold *p<*0.05 (right). RnR pools its top-3 undersampling-equivalent solutions per participant; each baseline contributes its single graph.

#### 4.3.3 The recovered connectome orients the canonical FNC picture

The significant edges form a coherent, literature-aligned circuit (Fig. 8a): a <u>sensory → visual hyperconnectivity hub</u> built on a hyperconnected somatosensory source. In SZ the directed edge from the postcentral gyrus component to the primary visual component is recovered more frequently than in controls (PoCG → calcarine, Δ *>* 0, the strongest edge in the map), and the same component is the source of the largest number of group-differing outgoing edges, to the caudate, superior temporal gyrus, anterior cingulate, thalamus and posterior cingulate (all Δ *>* 0; e.g. the thalamo-sensorimotor limb PoCG → Thal). These are component-level effective-connectivity relations, not monosynaptic anatomical projections. A somatosensory component that acts as a shared driver most likely reflects common sensory and motor drive rather than a direct postcentral influence on visual, striatal and cingulate cortex. These edges lie on the pairs that carry the most replicated resting-state finding in schizophrenia, thalamic and sensory hyperconnectivity to sensorimotor, auditory and visual cortex [45, 4, 15], and they add the orientation that an undirected measure could not supply. Our map encodes the thalamo-sensory limb of the canonical thalamocortical signature, but with a somatosensory source rather than a thalamic one: the thalamus appears as a target, and the elevated thalamo-sensorimotor coupling is oriented cortex-to-thalamus. The complementary association/prefrontal-hypoconnectivity limb requires prefrontal components that the tractable *N* =10 node set does not resolve, and is left to higher-resolution work (see Section 5). Figure 8b shows the spatial map of each component, making the distributed nature of these units explicit.

**Figure 8.**
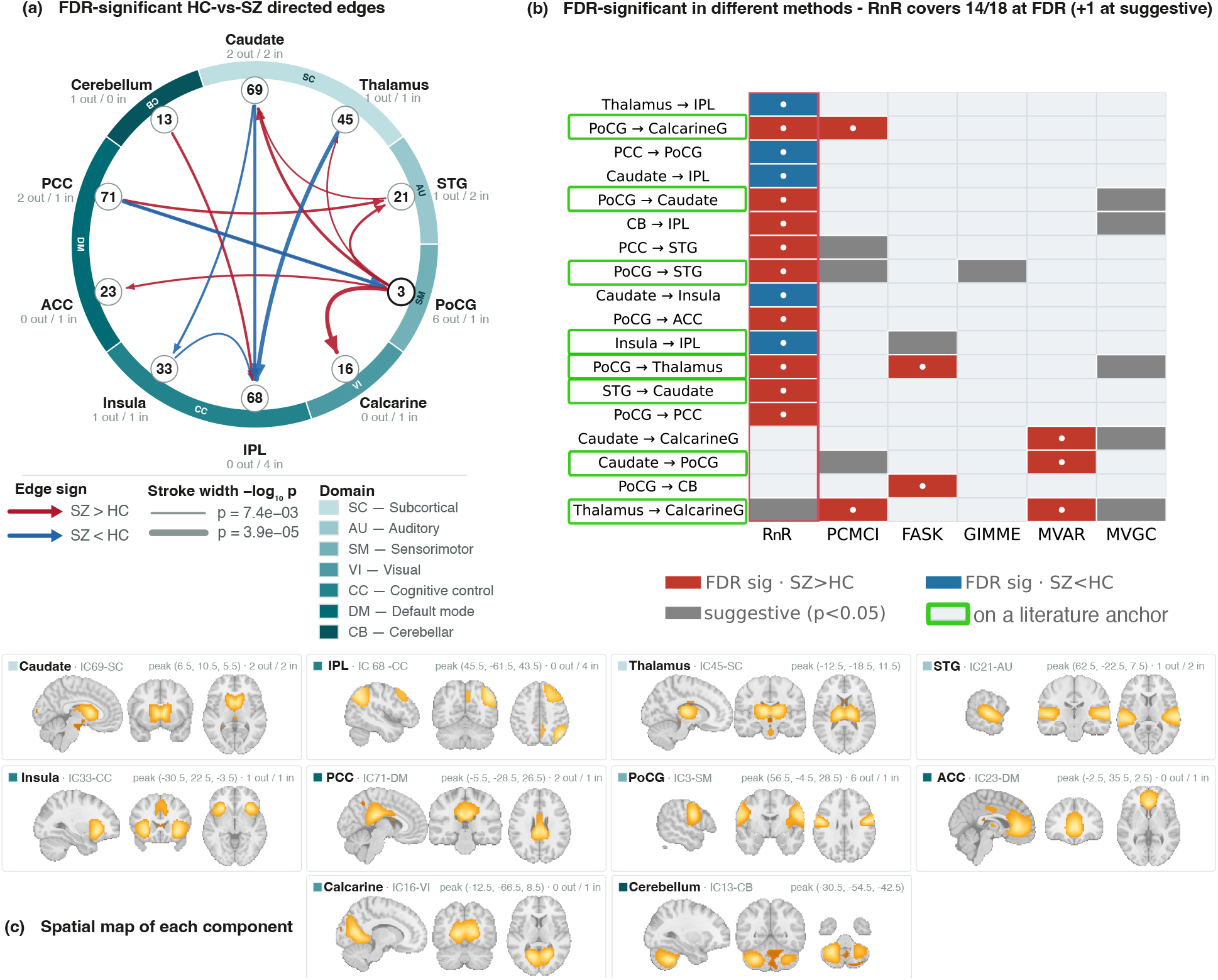
The recovered connectome and the spatial extent of its nodes. **(a)** The FDR-significant HC-vs-SZ directed edges of Fig. 8a, redrawn with the nodes ordered and coloured by NeuroMark functional domain. Arrows give causal direction, colour the sign of the SZ−HC effect (red SZ*>*HC, blue SZ*<*HC) and stroke width − log_10_ *p*. The number inside each node is its index in the NeuroMark template, and the bold outline marks the most connected node. **(b)** The group-ICA spatial map of each of the ten components, shown as sagittal, coronal and axial sections through that component’s published peak coordinate (positive tail, display threshold *z >* 5). Panel (b) makes explicit what a component label elides: each node is a distributed, weighted map rather than the structure at its peak. The postcentral component (IC 3) is a bilateral peri-central map even though its peak lies in the right hemisphere, and the inferior-parietal component (IC 68) extends over bilateral parietal and frontal cortex, whereas the thalamic (IC 45) and cerebellar (IC 13) components are compact and closely match their labels. A directed edge is therefore a relation between two components, not evidence of a projection between the structures at their peak coordinates (component selection in Section 4.3.1).

#### 4.3.4 RnR recovers most of the cross-method union

Across all methods, the union of FDR-significant edges contained 18 distinct edges (over *N* =10 nodes); RnR accounted for 14 of them at FDR, and a fifteenth, Thal → calcarine, at the suggestive level. Multiple method families converged on the sensory → visual hub: PoCG → calcarine was recovered by RnR and PCMCI, Thal → calcarine and Caudate → PoCG by MVAR, and PoCG → Thal by FASK. RnR’s output was not a strict superset of the others—MVAR uniquely contributed two striato-sensory edges—so the relationship is best described as convergence on a robust core plus a broad RnR-only periphery. This conclusion is consistent with RnR recovering the same structure as the single-graph methods, plus the edges visible only across timescales. Overlaying the curated literature anchors (green outlines in Fig. 8b) shows that RnR’s recovered edges include most of the displayed anchor pairs, so its larger yield is literature-aligned rather than indiscriminate.

Because RnR pools members of the undersampling equivalence class, its yield grows with pooling depth *k* (Fig. 9a). Literature recall (Eq. 4) rises in step with the edge count—from 0 at *k*=1 to 0.81 by *k*=10— indicating that deeper pooling adds edges that correspond to known SZ pairs rather than threshold noise. We report all primary results at a deliberately conservative *k*=3 (recall 0.375 at *N* =10). We are explicit that pooling inflates the effective sample to *k*(#participants) and that the edge test treats the 453*/*480 pooled graphs as independent when the *k* solutions of one participant are not. The conservative pooling depth and literature anchoring thus partially offset, but do not eliminate, this pseudo-replication. The reported *p*-values are conditional on the pooled-independence assumption; the strictly subject-level test (one graph per participant) is the conservative bound and yields no significant RnR edges for any method, so the differential set reported here should be read as the structure RnR surfaces under its native multiplicity, with Fig. 9a making the trade-off explicit.

**Figure 9.**
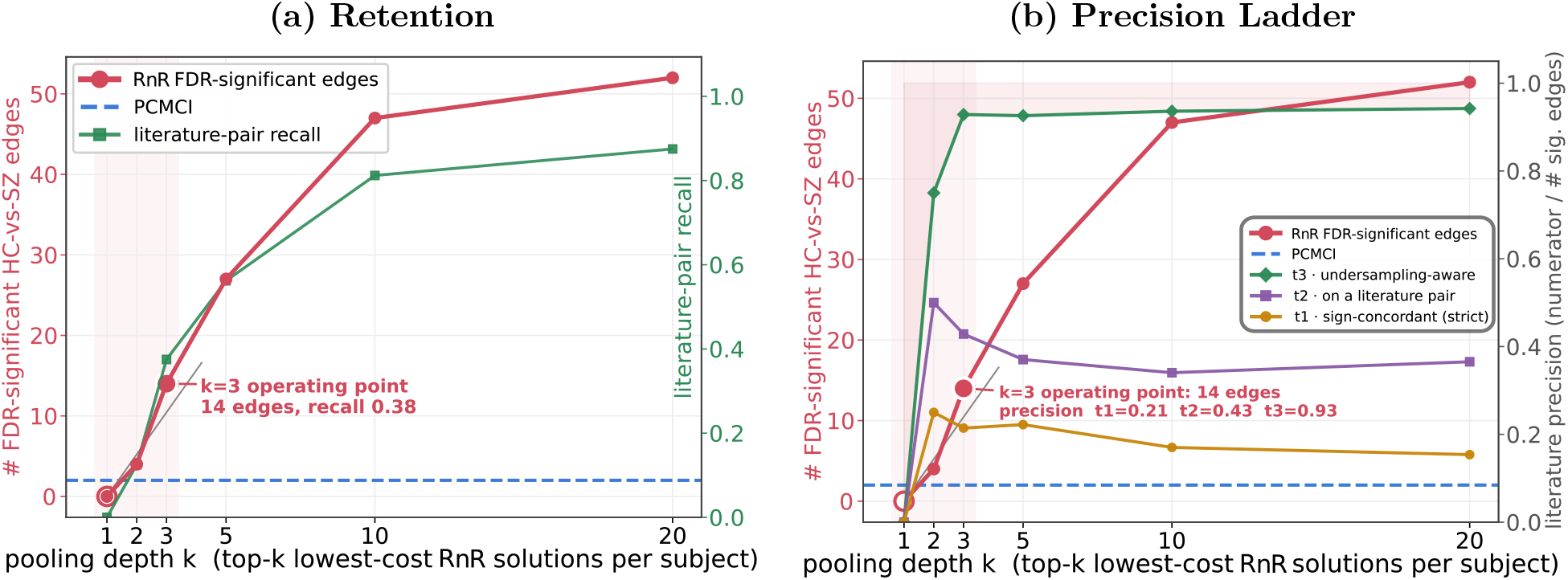
(a) Retention. RnR FDR-significant HC-vs-SZ edge count (red, left axis) and literature-pair recall (green, right axis) versus pooling depth *k*, with the single-timescale PCMCI baseline (blue dashed) for reference. Recall climbs in step with the edge count — deeper pooling adds literature-anchored structure rather than threshold noise — and we operate at the conservative *k*=3 (14 edges, recall 0.38). **(b) Precision ladder**. The three precision definitions versus *k* (right axis), with the edge count and PCMCI baseline for context (left axis): sign-concordant *π*_1_ (strict; “t1”), on-a-literature-pair *π*_2_ (direction-agnostic; “t2”), and undersampling-aware *π*_3_ (Eq. 7; “t3”), which credits novel directed edges and sign-discordant edges on fragile pairs. At the *k*=3 operating point *π*_1_=0.21, *π*_2_=0.43, *π*_3_=0.93; only the thin margin above *π*_3_ is a hard contradiction of a robust literature finding.

Precision tells a complementary story (Fig. 9b). Strict sign-concordant precision *π*_1_ is modest and declines with *k* (≈ 0.15–0.25). If we relax the hyper-/hypo-connectivity sign requirement (*π*_2_; justified because the undirected literature’s sign is a property of the net pair, not of a single directed limb) then precision reaches ≈ 0.3–0.5. Finally, the undersampling-aware *π*_3_ (Eq. 7)—crediting novel directed edges (on which the undirected literature is silent) and sign-discordant edges on fragile pairs (where undersampling can flip the apparent orientation)—reaches 0.93 at *N* =10. Only 1–3 edges are hard contradictions in which there is a wrong hyper-/hypo-connectivity sign on a pair whose orientation is timescale-robust, so undersampling cannot explain the mismatch. We interpret the undersampling-aware *π*_3_ as a permissive upper bound on <u>compatibility</u>—the fraction of edges that the literature gives no defensible reason to reject—rather than confirmed agreement. In particular, at *N* =10, the increase over *π*_2_ comes entirely from novel directed edges on which the undirected literature is silent, since no significant edge was classified undersampling-fragile at this resolution (see Section 4.3.5).

#### 4.3.5 Robust versus undersampling-fragile edge directions

Partitioning RnR’s pooled solutions by inferred rate (Eq. 6; 8,927 solutions at *N* =10, *u* ∈ {2, 3, 4, 5}) yielded a catalogue of 35 undersampling-robust directed pairs, those whose orientation is invariant across all observed voting rates (*u* ∈ {2, 3, 4, 5}), out of the 45 pairs at this resolution, and no undersampling-fragile pairs. Not a single pair’s dominant orientation flipped decisively with *u*: the other 10 pairs do change dominant orientation across voting levels, but never by a decisive margin (every level lies within 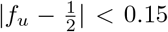 of a coin flip). Of the 14 FDR-significant edges, 11 were classified robust, including the sensory → visual hub, while the remaining 3 (Caudate–PoCG, Thalamus–PoCG, IPL–Insula) fall in that indecisive group and are left unclassified rather than fragile.

This absence of fragility is itself informative. At the coarse *N* =10 resolution, the inferred rates are low (dominated by *u*=2; see Section 3.1), so orientations are not yet being scrambled by temporal subsampling. Both undersampling theory [16] and our own simulations (in Section 5.1) predict that fragility, and the orientation reversals it produces, should emerge as spatial resolution sharpens toward the neural scale and the perceived rate climbs. Circuits whose directionality the undirected literature has long contested, notably the cortico-striatal-prefrontal arm, are where such reversals would be expected to surface first; resolving this requires the larger node sets discussed below (in Section 5) and is left to future work.

## 5 Discussion

Our central claim is methodological and conservative: applying undersampling-aware causal discovery to FBIRN reproduces the established functional-connectivity picture of schizophrenia where that picture is strong, expresses it in directed form, and adds a new, timescale-grounded interpretation only where the literature is itself unresolved. Three points support this reading.

RnR’s strongest and most reproducible edges involve the same pairs as those in the literature: thalamo-/sensory-cortical hyperconnectivity to auditory, somatomotor and visual systems [45, 4, 15], recovered here on the same source cohort (FBIRN phase-III) and group-ICA lineage used by **(author?)** [15]. This sub-cortical/sensory hyperconnectivity is the thalamo-sensory limb of the canonical thalamocortical signature [45, 4], encoded here as a single somatosensory source (postcentral gyrus), with the thalamus as a target rather than the driver. The complementary thalamo-prefrontal and cortico-striatal hypoconnectivity limb [4, 23] requires prefrontal components not resolved in our tractable node set and is left to higher-resolution work. Where our signs appear to disagree with a particular report, the disagreement falls exclusively on pairs that the literature itself reports as sign-unstable across static-vs-dynamic FNC [15], seed-vs-ICA methodology, illness stage (first-episode/early versus chronic; 34), and preprocessing. There is never disagreement on a high-confidence, directionally-determinate finding. (Consistent with this, cerebello-thalamo-cortical hyperconnectivity is reported as a stage-independent, trait-like signature; 12.)

Functional connectivity is, by construction, an <u>undirected</u> measure read at a <u>single</u> (slow) timescale. It therefore cannot state which region drives which, and it is exposed to the direction-reversal that temporal subsampling induces [16, 36, 22]. RnR addresses both shortcomings. It supplies orientation for couplings that have only been reported as magnitudes—e.g., it renders the postcentral–visual coupling elevated in SZ as a directed somatomotor → visual relation (PoCG → calcarine), an orientation no correlation-based study could supply—and it labels each orientation by its stability across undersampling rates. This reframes “novel” edges not as unsupported but as directed structure that the field’s dominant instrument cannot resolve, and reframes apparent sign disagreements as expected consequences of temporal subsampling— direction reversals induced by observing the process only at the coarser timescale—rather than as failures of replication. The same machinery yields a directed reading of network-level findings: a larger differential edge set reaching across functional domains is a directed correlate of the reduced network segregation reported graph-theoretically in SZ [26], and the directed insula → control/DMN edges speak to the triple-network “salience-switch” account [32, 42] in a form (effective, directed) not possible with the correlational evidence base.

The robust/fragile taxonomy presented here is a feature of the rate-agnostic analysis used here. It tells a downstream analyst which orientations remain stable across the evaluated rates, and which may change under different timescale assumptions in a single-timescale analysis. At the *N* =10 resolution, 11 of the 14 significant edges meet the robustness criterion, while three are unclassified. For instance, one of the robust edges is the sensory → visual hub. Therefore, the directed reading offered here is defensible, as it is stable across all undersampling rates we could evaluate. Finding few fragile orientations at this coarse resolution is consistent with the mechanism discussed in Section 5.1: heavy spatial aggregation leads to a low perceived undersampling rate, so theoretically possible orientation reversals [16] have not yet appeared. This same explanation also predicts, both from theory and simulation, that they will appear as resolution sharpens, and that circuits whose directionality the literature has long contested (notably the cortico-striatal-prefrontal arm) are where they should first surface. Testing these predictions will require the larger node sets discussed below.

### 5.1 Spatial resolution as the driver of the perceived undersampling rate

If temporal undersampling distorts causal structure as severely as we argue, it is fair to ask why two decades of single-timescale connectivity analyses have nonetheless produced a reproducible schizophrenia signature. We propose, and confirm in simulation, that a likely answer is spatial resolution. fMRI does not observe neurons, but rather spatial aggregates (voxels, regions, or independent components), each pooling millions of neurons whose fast <u>within</u>-aggregate interactions are never seen. Aggregation therefore potentially lowers the <u>perceived</u> undersampling rate—the rate one would infer from the aggregated signal—far below the true rate of the underlying neural process, because most fast causal steps are spent inside an aggregate and never cross between the observed nodes [25].

To quantify this possibility, we generated directed graphs at a realistic brain scale (*N* =53 nodes, the NeuroMark component count) organised into functional modules, undersampled them at a known true rate *u* (Eq. 1), and then coarsened them to progressively lower spatial resolutions by merging whole modules (see Fig. 11 for a coarsening example on a 16-node graph). As we move from per-node (*k*=53) down to two hemispheres (*k*=2), we determine the undersampling rate that would be perceived at each resolution. The rate compression is severe (Fig. 12a): a process undersampled at a true rate of 30 registers as *u* ≈ 19 at full per-node resolution but only *u* ≈ 7 once nodes are grouped into 12 functional-network modules (~ 4–5 components each, roughly the resolution of component-level fMRI) and as *u* = 1 at two-region resolution. The perceived rate falls monotonically as resolution coarsens and is bounded by the coarse graph’s own reachability (Fig. 12b); sparser, more physiologically realistic connectomes leave even more room for this compression. Figure 10 shows results for a single true rate (*u*=30): the perceived rate falls monotonically as spatial resolution coarsens and, at every resolution, is larger for sparser graphs.

**Figure 10.**
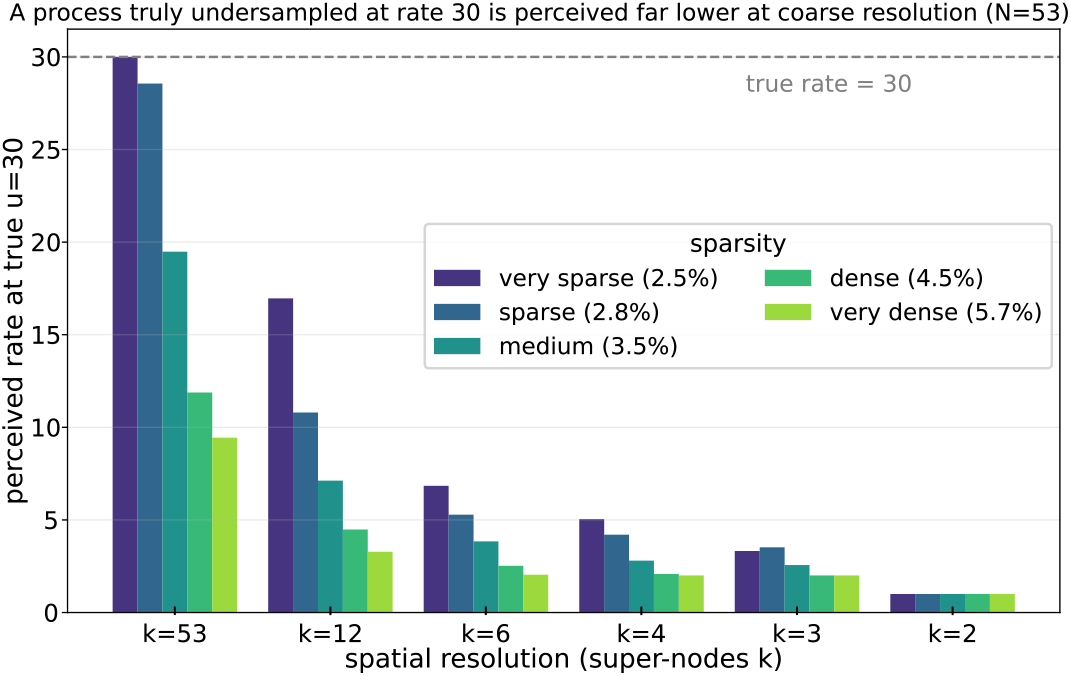
Spatial aggregation lowers the perceived undersampling rate (fixed true rate *u*=30). Mean perceived rate versus spatial resolution, grouped by graph sparsity (dashed line = the true rate); at full resolution only the sparsest graphs register rate 30 (denser graphs saturate first), and every coarsening drives it down to 1 at hemisphere resolution — density and coarsening compound. A process undersampled at a high true rate is perceived as only mildly undersampled once spatially aggregated; coarse spatial resolution suppresses the undersampling problem, higher resolution exposes it.

**Figure 11.**
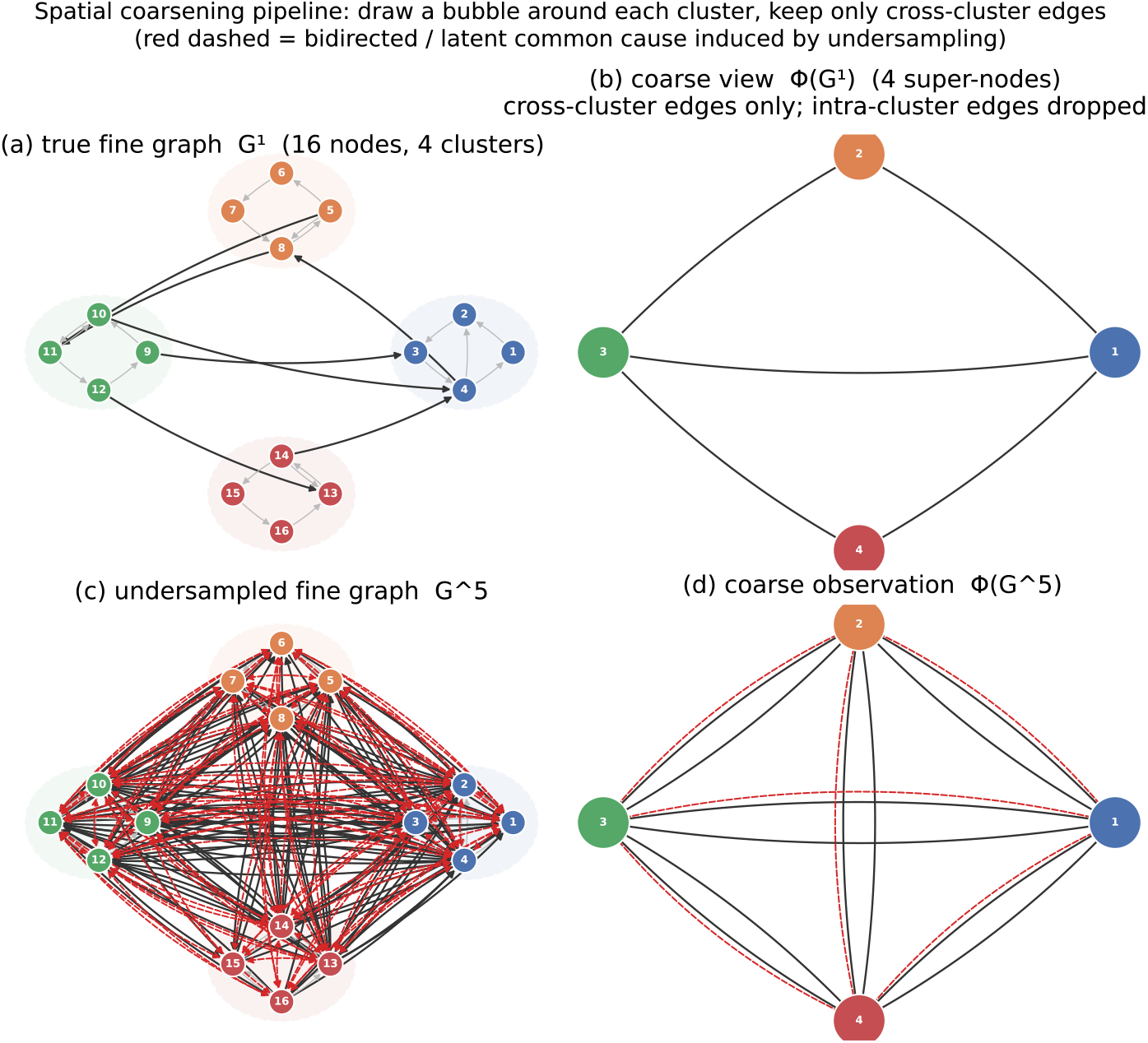
The spatial-coarsening operator (lowering spatial resolution). Shown on a 16-node, 4-cluster example. **(a)** the true fine graph, a bubble drawn around each cluster; **(b)** its coarse view Φ(*G*^1^) — one super-node per bubble, keeping only cross-cluster edges and dropping intra-cluster ones; **(c)** the fine graph undersampled to *G*^5^, already near-saturated (red dashed = bidirected/latent-common-cause edges induced by undersampling); **(d)** what the low-resolution observer sees, Φ(*G*^5^). Because coarsening and undersampling do not commute, the coarse observation saturates at a much smaller rate than the fine graph — the mechanism that lowers the perceived undersampling rate.

**Figure 12.**
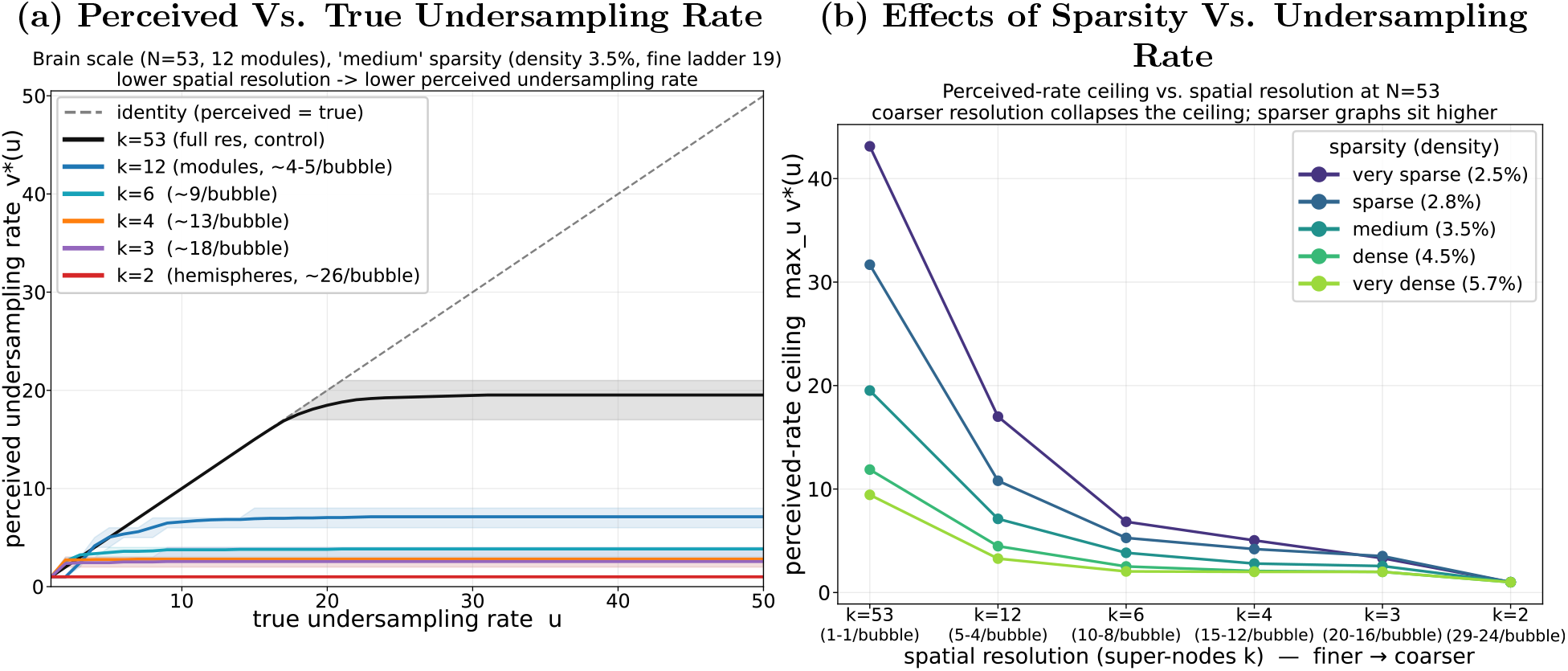
Spatial aggregation lowers the perceived undersampling rate. **(a)** Perceived versus true undersampling rate for simulated 53-node modular graphs (representative sparsity) as spatial resolution is coarsened by merging whole modules: full per-node resolution (*k*=53) tracks the true rate until the graph saturates, whereas every coarser resolution plateaus at a low ceiling (*k*=12 functional modules ≈ 7; *k*=2 hemispheres = 1). Dashed line: perceived = true. **(b)** Maximum perceived undersampling rate versus spatial resolution (super-nodes *k*; finer → coarser), one curve per graph sparsity (directed density 2.5–5.7%): the ceiling drops fastest at the first coarsening step and is highest for the sparsest — most physiologically realistic — graphs.

This explains why coarse-resolution analyses have been largely unaffected. At the coarse, whole-brain resolutions the field actually uses, the perceived undersampling rate should be small—consistent with the low rates RnR infers on FBIRN (*u* ≈ 2 for ~ 78% of solutions; see Section 3.1)—so a single-timescale estimator that implicitly assumes *u*=1 will be approximately correct and capable of recovering the robust undirected signature. At the coarse *N* =10 resolution in the FBIRN cohort, the inferred rate is low (*u* ≈ 2) and no recovered orientation is fragile (see Section 4.3.5): the directed reading is stable precisely because aggregation has held the perceived rate down. However, this seeming success is resolution-dependent, not reliable success. As spatial resolution improves, the single-timescale estimator should worsen: every step toward a finer, more neuron-like parcellation will raise the perceived rate and thereby risks of fragility and orientation reversals [16]. Even with coarse spatial regions, the perceived undersampling rate is strictly greater than one, leaving a residual undersampling distortion that a rate-aware method corrects and a single-timescale method inherits silently, pointing towards the need for a rate-aware method. As measurements and analyses move toward the true neural graph—the goal of effective-connectivity research—the field can less afford to ignore undersampling. Spatial aggregation has shielded the field so far, but as the field moves towards higher resolutions, we need to model undersampling explicitly, as RnR does.

### 5.2 Limitations

#### Shared sample with the reference FNC study

We analyse the same FBIRN phase-III cohort, pre-processing lineage and group-ICA template family as the canonical dynamic-FNC study of **(author?)** [15]. Because cohort composition, site mix and component definition are held fixed, differences between the directed connectome reported here and the published undirected one are attributable to the estimator rather than to the data, and the comparison in Section 4.3.3 is a concordance test on the same individuals. The design does not, however, provide an independent replication: whether the directed sensory-to-visual hub generalizes will require external cohorts such as COBRE, B-SNIP or the HCP Early Psychosis sample.

#### Illness stage and medication

The patient sample comprises predominantly chronic, clinically stable outpatients maintained on antipsychotic medication [24]. Chronicity and cumulative antipsychotic exposure both modulate the sign and magnitude of resting-state connectivity [40, 15] and neither is modeled here, so any part of the group difference attributable to treatment or illness duration is not separable in this design. Applying the same pipeline to first-episode and antipsychotic-naive samples would separate these effects and test whether the cortex-to-thalamus orientation of the thalamo-sensorimotor limb is present at illness onset.

#### Diagnostic specificity

Contrasting schizophrenia with healthy controls alone, we cannot establish that the recovered directed structure is specific to schizophrenia rather than common to psychosis. Work from the B-SNIP consortium has shown that resting-state connectivity abnormalities are substantially shared between schizophrenia and psychotic bipolar disorder [28, 27], and that biomarker-derived psychosis biotypes cut across DSM diagnostic boundaries [43, 14]. Settling the question requires the same analysis in a cohort containing schizoaffective and psychotic bipolar probands; until then the present results characterize this schizophrenia sample and cannot be claimed as diagnosis-specific.

#### Non-independence of the pooled solutions

The *k* solutions retained per participant are alternative members of a single undersampling equivalence class recovered from one time series, not independent observations, and the pooled test of Section 3.2 treats them as though they were, inflating the effective sample from 311 to 933. We operate at a deliberately conservative *k*=3 and state the subject-level bound explicitly: with one graph per participant, no method in the panel, RnR included, returns a significant edge. A test that respects the dependence structure would permute diagnostic labels at the level of the participant, carrying that participant’s *k* solutions as an inseparable block; we identify this as the appropriate exact test and leave it to work that also extends the node set.

#### Scope of the robustness classification and of the node set

“Robust” means invariant over the discrete rates realized by the solver (*u* ∈ {2, 3, 4, 5}, dominated by *u*=2) rather than over all conceivable *u*, and the sparser high-*u* strata carry the least support. The literature anchor encodes undirected magnitude signs that map onto, but are not identical with, our directed edge-presence effects, and component-to-region correspondences should be read against template peak coordinates before anatomical claims are drawn [17]. Finally, the node set is bounded by the computational cost of exact undersampling-aware recovery and omits circuits the literature also implicates, notably hippocampal, fuller cortico-striatal-limbic and motor components, and the prefrontal components needed to test the hypoconnectivity limb of the thalamocortical signature; extension to *N* =20 or *N* =53 is the natural next step, and is where theory predicts undersampling-fragile orientations should first appear.

## 6 acknowledgments

This work was supported by NIH R01MH129047 and in part by NSF 2112455. fBIRN data was supported by the National Center for Research Resources at the National Institutes of Health (NIH 1 U24 RR021992) and NIH 1 U24 RR025736.

## 7 Supporting Information

### 7.1 Clingo encoding and implementation of sRASL and RnR

This section connects the graph-theoretic definitions in Eqs. (1)–(2) to their computational implementation. The distinction between sRASL and RnR is important. Solver-based RASL (sRASL) encodes the noiseless inverse problem: it searches for causal-timescale graphs *G*^1^ and undersampling rates *u* for which the graph implied at the measured timescale is exactly the supplied graph ℋ[1]. Real-world noisy RASL (RnR) retains the same forward undersampling rules, but replaces exact agreement with weighted optimization because a graph estimated from finite fMRI data need not be the exact undersampled image of any *G*^1^ [2]. Both methods are implemented in the open-source `gunfolds` package [35] using the Answer Set Programming solver `clingo`.

#### 7.1.1 Why Answer Set Programming is appropriate

The recovery problem in Eq. (2) requires inversion of a many-to-one mapping. Given a causal-timescale graph *G*^1^ and an undersampling rate *u*, the forward operator in Eq. (1) determines the measured graph *G*^u^. In the reverse direction, however, a single measured graph can have multiple causal-timescale explanations. Recovery must therefore search over both the space of directed graphs and the possible undersampling rates.

For *n* nodes, there are *n*^2^ possible directed edges when self-connections are allowed and consequently 2^n^ possible causal-timescale graphs. A procedural algorithm must generate candidates, calculate their undersampled graphs, compare each result with ℋ, and implement specialized pruning rules. Adding new structural information then requires modifying the search procedure itself.

Answer Set Programming (ASP) provides a natural alternative because it is declarative. Rather than specifying how to traverse the candidate space, the program specifies what constitutes a candidate and what conditions make it valid. The solver is responsible for finding the objects that satisfy those conditions. The forward undersampling operator, agreement with the observed graph, strongly connected component restrictions, the density prior, and the edge-evidence weights can therefore be expressed as separate, composable logical statements.

This representation is particularly appropriate here because the scientific answer is generally a set of graphs rather than a unique point estimate. In exact sRASL, the answer sets returned by the solver correspond to the members of the equivalence class in Eq. (2), within the represented rate domain and modelling assumptions. RnR uses the same representation but ranks feasible explanations according to their disagreement with a noisy estimated graph.

#### 7.1.2 The ASP language used in the encoding

An ASP program is commonly organized according to a <u>generate–define–test</u> structure. Facts describe the particular problem instance. Choice rules generate candidate objects. Ordinary rules derive the consequences of each candidate. Integrity constraints eliminate candidates that violate the model. In RnR, weak constraints replace strict rejection with weighted penalties.

The principal ASP constructs used in the implementation are as follows:

- **Facts** are unconditional statements describing the problem instance. For example, the measured directed edge 1 → 3 can be represented by `hdirected(1,3).`, while `hbidirected(2,7).` Represents 2 ↔ 7 in ℋ. A fact such as `scc(1,2).` assigns node 1 to strongly connected component 2. The shorthand `node(1..n).` produces one fact for every node.
- **Ordinary rules** have the form `Head :- Body.`. The rule means that the head must hold whenever the body holds. Variables begin with uppercase letters and are instantiated over their finite domains during grounding. For example,

~~~
directed(X,Y,1) :- edge1(X,Y).
~~~

states that a candidate edge from *X* to *Y* is a directed walk of length one.
- **Choice rules** generate the candidate space. The rule

~~~
{edge1(X,Y)} :- node(X), node(Y).
~~~

allows each possible causal-timescale edge to be present or absent. Cardinality bounds can restrict the number of selected atoms. The rule

~~~
1 {u(1..maxu)} 1.
~~~ selects exactly one undersampling rate for each answer set.
- **Integrity constraints** have an empty head and take the form `:- Body.`. A candidate is rejected whenever the body is true. Exact sRASL uses integrity constraints to require the graph derived at the selected rate to equal ℋ.
- **Weak constraints** have the form

~~~
:~ Body. [W@P,…]
~~~

and do not reject a candidate. Instead, a satisfied weak constraint adds weight *W* to the candidate’s objective at priority *P*. `clingo` minimizes higher priority levels before lower ones. RnR uses weak constraints to penalize disagreements with noisy or uncertain features of ℋ. The central graph predicates have direct interpretations:
- `edge1(X,Y)` denotes *X* → *Y* in the unknown causal-timescale graph G^1^.
- `u(U)` denotes the candidate undersampling rate.
- `directed(X,Y,L)` states that G^1^ contains a directed walk of length *L* from *X* to *Y*. At *L* = *u*, this implies *X* → *Y* in G^u^.
- `bidirected(X,Y,U)` states that *X* and *Y* have a common ancestor that reaches both through directed walks of the same length *L < U*. This implies *X* ↔ *Y* in G^U^.
- Predicates beginning with `h`, such as `hdirected` and `hbidirected`, describe the measured input graph ℋ rather than a candidate causal graph.

After grounding the variables over the finite node and rate domains, `clingo` searches the resulting propositional program for stable models, or “answer sets.” In this application, the displayed atoms in one answer set encode one candidate graph–rate pair (G^1^, *u*).

#### 7.1.3 Generating a candidate causal graph

The following ASP fragment generates the space of candidate causal-timescale graphs and possible under-sampling rates. The constant `maxu` is the maximum rate considered by the solver.

~~~
#const n = 10.
#const maxu = 20.
node(1..n).
1 {u(1..maxu)} 1.
{edge1(X,Y)} :- node(X), node(Y).
~~~

The first choice rule selects exactly one rate. The second independently selects the presence or absence of every possible directed edge in *G*^1^. Consequently, the rule represents the full candidate space without requiring a manually written loop over graphs.

Self-connections are included because an edge *X*^t−1^ → *X*^t^ represents temporal persistence of the same variable and can affect both directed reachability and SCC stability.

#### 7.1.4 Encoding the forward undersampling operator

The following rules derive the directed and bidirected edges that a candidate *G*^1^ would produce after under-sampling. They constitute a direct discrete encoding of Eq. (1).

~~~
directed(X,Y,1) :-
    edge1(X,Y).
directed(X,Y,L) :-
    directed(X,Z,L-1),
    edge1(Z,Y),
    L <= U,
    u(U).
bidirected(X,Y,U) :-
    directed(Z,X,L),
    directed(Z,Y,L),
    node(X;Y;Z),
    X < Y,
    L < U,
    u(U).
~~~

The first rule establishes the base case: every candidate edge is a directed walk of length one. The second rule recursively extends directed walks. If *X* reaches *Z* in *L* − 1 steps and *Z* → *Y* is an edge of *G*^1^, then *X* reaches *Y* in *L* steps. Therefore, `directed(X,Y,U)` holds exactly when the candidate graph induces *X* → *Y* at the selected rate *U*.

The final rule implements the hidden-common-cause component of undersampling. It derives a bidirected edge between *X* and *Y* when some node *Z* reaches both through directed walks of equal length *L < U*. The common cause lies between measured time points and is therefore unobserved at the measured timescale. The restriction *X < Y* stores each unordered bidirected pair only once.

These rules are shared by exact sRASL and RnR. The difference between the methods lies primarily in how the derived graph is compared with ℋ.

#### 7.1.5 Exact matching in sRASL

For an error-free measured graph, sRASL enforces equality between the derived graph *G*^u^ and ℋ. The following four integrity constraints exclude, respectively, an extra directed edge, an extra bidirected edge, a missing directed edge, and a missing bidirected edge:

~~~
:- directed(X,Y,U),
not hdirected(X,Y),
u(U).
:- bidirected(X,Y,U),
not hbidirected(X,Y),
X < Y,
u(U).
:- not directed(X,Y,U),
hdirected(X,Y),
u(U).
:- not bidirected(X,Y,U),
hbidirected(X,Y),
X < Y,
u(U).
~~~

The first two constraints prevent the candidate from producing edges that are absent from ℋ. The final two prevent the candidate from omitting edges that are present in ℋ. Together they impose

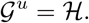

Every answer set that survives these constraints is therefore a valid pre-image of ℋ under the undersampling operator, within the finite rate domain and the assumed model class.

This exactness is also a limitation when ℋ is estimated from finite data. Statistical noise can add, omit, or misorient edges, and only a subset of mixed directed–bidirected graphs can be generated by undersampling a purely directed *G*^1^. An estimated graph may consequently have no exact pre-image, in which case the exact ASP program is unsatisfiable.

#### 7.1.6 Strongly connected component constraints

sRASL improves computational efficiency by encoding structural information about strongly connected components (SCCs). Facts of the form `scc(X,K)` assign node *X* to component *K*, while facts such as `dag(K,L)` record permitted directions between components. A representative SCC restriction is

~~~
:- edge1(X,Y),
scc(X,K),
scc(Y,L),
K != L,
sccsize(L,S),
S > 1,
not dag(K,L).
~~~

Cycles remain permitted within an SCC, but connections between different SCCs must be compatible with the component-level directed acyclic graph. Under the loop-length condition stated in the Methods, SCC membership is stable under undersampling. Supplying the SCC facts before search therefore eliminates large families of structurally incompatible candidates.

These restrictions are one reason sRASL is substantially faster than search-based RASL. Instead of generating a graph and discovering only after undersampling that its component structure is impossible, the ASP solver incorporates the structural restrictions during search. The present analysis uses the domain-constrained SCC encoding described in Section 3.1.

#### 7.1.7 The finite rate domain

The executable program searches over a finite range of rates controlled by `maxu`. This is both a modelling decision and a computational necessity. Repeatedly undersampling a finite graph does not produce new graphs indefinitely. Under the same loop-length condition that stabilizes SCC membership, the sequence eventually reaches a repeated or stable state [36].

The implementation can therefore suppress later occurrences after the compressed graph repeats an earlier state. Exact enumeration is complete relative to the represented rate range and the stated structural assumptions. The upper bound should be selected using domain knowledge about the relationship between the measurement interval and the plausible causal timescale. It is not itself learned continuously from the data; rather, the solver selects a discrete rate within the specified range.

#### 7.1.8 RnR as a weighted relaxation of sRASL

RnR retains the graph generator and forward undersampling rules, but replaces exact matching with weighted optimization. The four hard constraints are converted into weak constraints:

~~~
:~ directed(X,Y,U),
no_hdirected(X,Y,W),
u(U).
[W@0,X,Y,1]
:~ bidirected(X,Y,U),
no_hbidirected(X,Y,W),
X < Y,
u(U). [W@1,X,Y,2]
:~ not directed(X,Y,U),
hdirected(X,Y,W),
u(U).
[W@0,X,Y,3]
:~ not bidirected(X,Y,U),
hbidirected(X,Y,W),
X < Y,
u(U).
[W@1,X,Y,4]
~~~

An `h*` fact records an edge observed in ℋ, while a `no h*` fact records an observed absence. The weight *W* is a non-negative integer derived from the strength of the upstream evidence. The precise priority levels are configurable; the schematic levels above separate bidirected-structure decisions from directed orientation decisions.

The first two weak constraints penalize edges produced by the candidate that are absent from ℋ. The last two penalize edges present in ℋ but missing from the candidate’s measured-scale image. Unlike an integrity constraint, a weak constraint does not reject the candidate. It records the disagreement as part of its cost.

Strongly supported observed edges receive large omission penalties and are therefore expensive to remove. Weakly supported edges receive smaller penalties and can be removed if doing so yields a more structurally coherent explanation. Conversely, adding an unsupported edge ordinarily receives a high penalty. The resulting objective is a weighted minimum-change problem rather than an exact graph-matching problem.

The trailing numeric field in each cost tuple is a type tag. It distinguishes directed and bidirected mismatch classes associated with the same node pair. Without distinct tags, two ground weak constraints with identical weights, priorities, and node terms can be treated as the same objective element and counted only once.

#### 7.1.9 Evidence weights

Let *A* denote the lagged association matrix returned by the upstream estimator and let *s*_*ij*_ ∈ [0, 1] be the normalized evidence magnitude for the ordered pair (*i, j*). A representative weighting scheme is

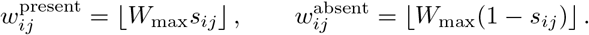

The first quantity is the cost of omitting an edge that the upstream method selected. The second is the cost of adding an edge that it did not select. A selected edge with strong evidence is therefore expensive to omit, while a rejected edge with very weak evidence is expensive to add.

In the present pipeline, the lagged and contemporaneous evidence matrices are obtained from the PCMCI output [38]. The same ASP formulation can accept scores from other first-order estimators, including Granger-causal or multivariate autoregressive methods. Prior anatomical information can also be incorporated by increasing or decreasing the relevant edge-level penalties.

#### 7.1.10 Density regularization

RnR additionally discourages candidate graphs whose density is incompatible with the expected sparsity of brain networks. A simplified encoding is

~~~
edge_count(C) :-
    C = #count {X,Y : edge1(X,Y)}.
density_error(E) :-
   edge_count(C),
   target_edges(T),
   E = |C-T|.
:~ density_error(E).
[E@2,E]
~~~

The first rule counts the directed edges in the candidate *G*^1^. The second computes the absolute deviation from the target edge count. The weak constraint penalizes that deviation.

This term is a regularizer, not a statement that the true density is known. Its role is to prevent noisy edge evidence from driving the solution toward nearly empty or nearly complete graphs. When used with a hard admissible density window, the implementation can widen the window or fall back to a purely soft penalty if the initial bounds make the instance unsatisfiable.

The cost can therefore be written schematically as

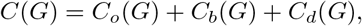

where *C*_o_ measures disagreement with the density target, *C*_b_ measures bidirected-edge discrepancies, and *C*_d_ measures directed-edge discrepancies. Priority levels implement a coarse-to-fine ordering, while the evidence weights resolve trade-offs within each level.

#### 7.1.11 Handling mutual directed pairs

RnR can operate as a meta-solver on the result of an existing causal discovery method. An upstream method without a bidirected-edge primitive may represent dependence due to an undersampling-induced hidden common cause as a mutual directed pair, *i* → *j* and *j* → *i*.

For such a pair, RnR can offer *i* ↔ *j* as an additional low-penalty explanation. The solver is not forced to choose the bidirected interpretation. Depending on the remaining structural constraints and evidence weights, it can retain one direction, retain both directions, choose the bidirected edge, or choose a compatible combination.

In the PCMCI front end, contemporaneous links can be represented directly as bidirected edges in the compressed graph supplied to RnR. Both mechanisms serve the same conceptual purpose: they prevent the representational limitations of the upstream estimator from being treated as exact knowledge about the causal-timescale graph.

#### 7.1.12 Solver execution and answer-set decoding

For each participant, the implementation proceeds as follows:

1. PCMCI, using the settings reported in Section 3.1, estimates a lag-one graph and edge-level evidence from the participant’s component time series.
2. The PCMCI result is converted to a compressed measured graph ℋ. Directed and bidirected presence and absence facts, their integer weights, the SCC-domain facts, the density prior, and the finite rate domain are added to the ASP program.
3. `clingo` grounds the program and searches jointly over the possible `edge1/2` atoms and the undersampling rate. In exact sRASL mode, an emitted answer set represents a graph–rate pair that satisfies G^u^ = ℋ. In RnR mode, the returned models are accompanied by their objective costs.
4. Output is restricted to the predicates carrying the result:

~~~
#show edge1/2.
#show u/1.
~~~

Intermediate atoms used to derive paths, bidirected edges, SCC restrictions, and density are omitted from the displayed answer set.
5. The shown `edge1(X,Y)` atoms are decoded as the directed edges of one causal-timescale graph. The shown `u(U)` atom provides its inferred rate. The Python layer converts these atoms into the internal `gunfolds` graph representation and associates the model with its objective cost.
6. Candidate models are ordered by cost. Let *c*_min_ be the lowest cost found for that participant. RnR defines a working candidate pool using the relative criterion

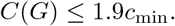

The *k* = 3 lowest-cost retained members are used in the group analysis, or all retained members if fewer than three are available.

A displayed answer set can therefore take the form

~~~
edge1(1,1) edge1(1,3) edge1(3,7) edge1(7,2) u(2)
~~~

which represents a causal-timescale graph containing the four displayed edges and an inferred undersampling rate of two.

#### 7.1.13 Interpretation of the returned set

The outputs of exact sRASL and RnR have related but distinct meanings. In exact sRASL, the returned answer sets form an equivalence class defined by exact equality to ℋ, within the represented rate range and modelling assumptions. Every member is a distinct causal-timescale explanation of the same measured graph.

In RnR, ℋ is treated as an uncertain estimate. The retained set is therefore a cost-bounded approximation to the exact equivalence class. Every candidate obeys the forward undersampling and structural rules, but candidates may disagree with uncertain features of ℋ at an explicitly recorded cost. A low cost means that relatively little well-supported information had to be changed; it does not prove that the candidate is the unique causal graph.

For this reason, the downstream analysis does not interpret a single answer set as the uniquely identified neural network. Instead, it retains several low-cost solutions and analyzes their shared and variable features. Edge frequency across solutions measures how consistently an edge is supported within the recovered candidate set. Partitioning the solutions by inferred *u* additionally makes it possible to determine whether a dominant orientation remains stable across rates or changes when a different temporal relationship between the measured and causal processes is assumed.

This solution-set interpretation is the central connection between the ASP implementation and the empirical analyses in the main text. The solver makes the ambiguity induced by temporal undersampling explicit, while the subsequent edge-frequency and robustness analyses determine which features of that ambiguity are scientifically stable enough to report.

## 8 Technical Terms

**Effective connectivity** — the directed, causal influence one brain region exerts over another, as opposed to mere statistical correlation.

**Functional connectivity** — statistical dependence (typically correlation) between the activity time series of brain regions; undirected.

**Temporal undersampling** — sampling a process more slowly than its own dynamics, so several fast steps collapse into one measured step.

**BOLD signal** — the blood-oxygenation-level-dependent fMRI signal, an indirect, slow proxy for underlying neural activity.

**Equivalence class** — the set of distinct causal graphs that are all equally consistent with the same observed data.

**Rate-agnostic structure learning** — causal discovery that does not assume the sampling rate matches the causal timescale.

**Answer Set Programming** — a declarative logic-based method for solving constraint-satisfaction and optimization problems.

**Independent component analysis** — a data-driven method that separates fMRI data into spatially independent networks (components).

**False discovery rate** — the expected proportion of false positives among results declared significant; controlled here by Benjamini–Hochberg.

**Undersampling-robust / -fragile orientation** — an edge direction that stays fixed across inferred sampling rates (robust) versus one that flips (fragile).

